# Comparative transcriptomic validation of candidate geroprotectors in human iPSC-derived neurons

**DOI:** 10.64898/2026.11.23.768196

**Authors:** Elise M. MacDonald, Natalie Lloyd, Nicholas J. Skvir, Carlos Villacorta-Martin, Samuel F. Cranston, Thomas Girke, Nicholas J. Schork, Paola Sebastiani, George J. Murphy

## Abstract

The discovery of geroprotectors, defined as interventions that slow aging-related deterioration and preserve function, remains constrained by reliance on lifespan studies. Lifespan studies take years, are costly and resource intensive, and return a whole-organism endpoint dominated in mice by neoplastic death. These studies cannot be readily scaled and do not specify which tissue(s) are protected by geroprotectors. Human induced pluripotent stem cell (iPSC)-derived cells provide a scalable alternative that enables defined lineages and genetically diverse donors to be studied in parallel. However, reprogramming resets epigenetic age, and whether iPSC-derived cells can report responses relevant to aging has remained unclear. Here, we evaluated eight candidate geroprotectors in iPSC-derived neurons using multiple classes of aging-associated transcriptomic readouts. Aging-associated pathways, including mTORC1 and MEK-ERK signaling, were intact and pharmacologically tunable in these cells. Genetic background accounted for most transcriptional variance, with compound responses differing in magnitude and reproducibility between donors. PI-103 suppressed the mTORC1 signature and increased DCX and BDNF expression, while also inducing genes associated with older age in two independent human excitatory-neuron references. These findings indicate that distinct aging measures capture different dimensions of drug response and that no single readout is sufficient to define geroprotection in our model. Together, these results support iPSC-derived cells as a scalable screening tool for candidate geroprotectors, enabled by the transcriptional aging readouts now available, with the potential to resolve tissue- and donor-dependent responses as additional lineages and donors are incorporated.

## Introduction

Aging is the dominant risk factor for most chronic diseases, including neurodegeneration, cancer, and cardiovascular disease (Kennedy *et al*., 2014; López-Otín *et al*., 2023). This observation underpins the geroscience hypothesis: interventions that slow the underlying rate of biological aging should delay the onset of many age-related diseases at once, rather than one at a time. Compounds pursued on this premise are termed geroprotectors, broadly defined as substances that slow aging, repair age-associated damage, or extend healthy lifespan (healthspan) (Moskalev *et al*., 2016). More than 200 compounds have been reported to meet this description in organisms ranging from yeast to rodents (Moskalev *et al*., 2015), yet none is approved clinically for the treatment of aging itself.

Notably, geroprotector discovery has a throughput bottleneck. Lifespan studies in mice take three to four years, are expensive, and yield a single endpoint. The Interventions Testing Program (ITP) has evaluated 54 compounds in two decades of operation in over 30,000 mice (Nadon *et al*., 2017; Jiang *et al*., 2025). To date, the ITP has identified 13 compounds that significantly extend lifespan, 12 of which only or preferentially work in male mice (Harrison *et al*., 2009; Garratt *et al*., 2017). Additionally, approximately 80% of UM-HET3 mice die of neoplastic disease, so the endpoint is dominated by one mechanism (Lipman *et al*., 2004). Under these criteria, a compound that postpones lethal cancer scores as a geroprotector regardless of what it does to the brain, or any other organ systems. Importantly, a lifespan study reports that an animal lived longer, not which tissue(s) was protected or through what mechanism.

Human induced pluripotent stem cell (iPSC)-derived cells address many of these constraints. iPSCs are expandable without limit, so a screen is not bounded by material. They can be derived from as many donors as a study design requires, making genetic diversity an essential variable rather than a constraint. They can be reproducibly directed to defined lineages, and the resulting tissue is human-specific as opposed to coming from a model organism, enhancing human translatability. Readouts are obtained in weeks rather than years. Their use in aging research is nevertheless limited by a fundamental paradox: the same reprogramming process that creates the model removes much of the donor’s age-associated epigenetic state and differentiated progeny retain immature features (Takahashi *et al*., 2007; Olova *et al*., 2019). This raises the question: What does it mean to call a compound a geroprotector in a system that does not have an epigenetic age?

Recent work changes what is possible, as classes of transcriptional aging readouts have emerged. Classical epigenetic biological aging clocks, such as the Horvath Clock (Horvath, 2013) and PhenoAge (Levine *et al*., 2018), are not useful tools in the context of iPSC-derived tissues. Due to the reprogramming process, these clocks consistently report ages near zero in iPSC-derived lineages (Kabacik *et al*., 2022; Skvir *et al*., 2026). Transcriptomic clocks estimate relative transcriptional age or mortality risk from transcriptomic expression alone (Tyshkovskiy *et al*., 2026). Aging rate indicators (ARIs) are another class of aging readouts. ARIs are defined as measurable features that discriminate slow-aging animals from normal aging animals and thus carry no information about biological age by design (Miller, Li and Garcia, 2023). Previous studies identified twelve candidate ARIs, shared across at least ten varieties of slow-aging mice, including reduced mTORC1 activity, reduced MEK1/ERK signaling, and increased BDNF and DCX in the hippocampus (Erickson *et al*., 2010; Austad, Kaeberlein and Miller, 2026). Here, we argue that ARIs and transcriptomic clocks are potential readouts for geroprotector identification in iPSC-derived systems.

Interestingly, the ARI framework also anticipates tissue specificity. Interventions may slow age-related change in particular organ systems rather than uniformly. One clear exception to this uniformity has been documented: rapamycin fails to raise UCP1 in white adipose tissue while affecting every other tested indicator (Li *et al*., 2023). As such, indicators are assigned to specific tissues. This raises the important distinction that if geroprotection is tissue-specific, screening for geroprotective compounds must be done in the tissue of interest. Excitingly, that is precisely what an iPSC model provides and what primary human tissue cannot deliver at scale. iPSC models offer a route to such comparisons, because the same donor lines can be differentiated into multiple lineages and screened in parallel. The present study evaluates a single lineage, cortical neurons, as a first step toward these cross-lineage comparisons.

Here, we test whether transcriptional readouts can be used to evaluate candidate geroprotectors in human iPSC-derived neurons. Compounds were selected by strength of prior evidence: rapamycin and trametinib as established model organism geroprotectors, PI-103 as a mechanistically related comparator without direct lifespan evidence, and the remaining five as exploratory candidates (**Table 1**). Ultimately, we aim to test if these cells report compound pharmacology faithfully, if ARIs are expressed and drug-responsive in them, whether weaker responses are coordinated across indicators, and ultimately whether any compounds stand out as candidate geroprotectors. We then applied human brain aging signatures to the same data, to test whether biomarker-type readouts add information to the present geroprotector screen in an iPSC system.

**Table 1.** Candidate geroprotective compounds tested in iPSC-derived cortical neurons. Mechanism, predicted Hallmark pathway effects, and prior aging-related evidence for each of the eight compounds. ↑, predicted enrichment; ↓, predicted depletion.

| Compound | Class / primary target | Expected pathway effect | Prior aging evidence |
|---|---|---|---|
| Rutin (RUT) | Flavonoid glycoside (quercetin-3-O-rutinoside); antioxidant and anti-inflammatory | ↓ Inflammatory response<br>↓ Reactive oxygen species pathway<br>↓ TNFα signaling via NF-κB | Extended lifespan in *Drosophila* in a hormetic, dose-dependent manner (Chattopadhyay *et al.*, 2017). |
| Sitagliptin (SIT) | Dipeptidyl peptidase-4 (DPP-4) inhibitor | ↓ IL6–JAK–STAT3 signaling<br>↓ Inflammatory response<br>↓ TNFα signaling via NF-κB | Extended lifespan in *C. elegans* via insulin/IGF-1 and dietary restriction-like signaling (Ye *et al.*, 2023). |
| Genistein (GEN) | Soy isoflavone; estrogen receptor agonist and tyrosine kinase inhibitor | ↑ Estrogen response (early) | Extended lifespan and stress resistance in *C. elegans* (Lee *et al.*, 2015). |
| Ebselen (EBS) | Organoselenium glutathione peroxidase mimetic; antioxidant | ↑ Oxidative phosphorylation<br>↓ Reactive oxygen species pathway<br>↓ Unfolded protein response | Raised GSH:GSSG ratio, reduced oxidative DNA damage, and increased cumulative population doublings of human CD4+ T-cell clones *in vitro* (Marthandan *et al.*, 2013). |
| Calpeptin (CAL) | Cell-permeable calpain inhibitor | ↓ Epithelial–mesenchymal transition | Genetic calpain inhibition reduced kidney and vascular inflammaging and senescence markers in aged mice; lifespan was not extended (Hanouna *et al.*, 2017). |
| PI-103 (PI) | Dual class I PI3K/mTOR inhibitor | ↓ mTORC1 signaling<br>↓ PI3K–AKT–mTOR signaling<br>↓ Apoptosis | Partial genetic inactivation of PI3K p110α extended lifespan in male mice (Foukas *et al.*, 2013). |
| Trametinib (TRA) | Allosteric MEK1/2 inhibitor | ↓ KRAS signaling<br>↓ MYC targets<br>↓ TNFα signaling via NF-κB | Extended lifespan in *Drosophila* and mice via inhibition of Ras–ERK–ETS signaling (Slack *et al.*, 2015; Gkioni *et al.*, 2025)11/20/26 4:01:00 PM |
| Rapamycin (RAP) | Allosteric mTORC1 inhibitor | ↓ mTORC1 signaling | Extended lifespan in genetically heterogeneous mice, including when started late in life (Miller *et al.*, 2011) |

## Methods

### iPSC culture and neuronal differentiation

iPSC lines used in this study were generated and differentiated into neurons as described in (Dowrey *et al*., 2025). Briefly, iPSCs were engineered to have a tetracycline-inducible neurogenin 2 (NGN2) cassette (Addgene 172115) using a PiggyBac plasmid EF1α-transposase (Addgene 172116) as described previously (Fernandopulle *et al*., 2018). Three days of tetracycline induction generated neural progenitor cells. Following induction, NPCs were replated for further maturation for 14 days.

### Dosing with compounds

Compounds and catalogue numbers are listed in **Table S1**. Stocks were prepared in DMSO and diluted through an intermediate before addition to medium. Neurons were treated for 48 hours (n=4 per compound, with 2 donors across 2 independent differentiations) at the final concentrations given in **Table S1**.

### RNA extraction, sequencing, and QC

RNA was isolated from cells using the Qiagen AllPrep DNA/RNA kit (Qiagen, 80284). Poly(A)-enriched mRNA libraries were sequenced by Novogene Co. (Sacramento, CA) on an Illumina NovaSeq 6000 S4 flow cell (paired-end 150 bp), with a minimum of 20 million reads per sample. Read quality was assessed with FastQC. Reads were aligned to the GRCh38 primary assembly with STAR 2.7.10b using the Ensembl release 92 annotation, and gene-level counts were generated with featureCounts (Subread 2.0.3) in paired-end mode. Genes with zero counts across all samples were removed, and counts were converted to log2 counts per million (CPM). Quality assessment included evaluation of library size distributions and principal component analysis (PCA).

### Differential expression and GSEA

Differential expression was tested with limma-trend (empirical Bayes moderation with an intensity-dependent prior and robust hyperparameter estimation) on log2 CPM values. The pooled model was ∼ 0 + treatment + cell_line, giving each compound a contrast against vehicle while blocking on genetic background. Per-line effects used ∼ 0 + line:treatment with within-line contrasts against that line’s vehicle. Differentially expressed genes were called at Benjamini-Hochberg adjusted p < 0.05 within each contrast. Cross-line reproducibility was assessed by selecting the 500 most significant genes in one line and correlating their effect estimates with the other line (Spearman).

MEK-ERK activity was scored using the 10-gene MAPK Pathway Activity Score (Wagle *et al*., 2018). mTORC1 activity was scored using the MSigDB Hallmark MTORC1_SIGNALING gene set. Scores are the mean z-score across signature genes, computed across all samples; treatment effects were tested with lm(score ∼ treatment + cell_line) and Benjamini-Hochberg corrected across treatments. Gene set enrichment used fgsea on limma moderated t-statistics, against the 50 MSigDB Hallmark sets, with correction applied across the full Hallmark collection within each contrast. Expected pathway directions were specified in advance from each compound’s known mechanism (**Table 1**) and results are reported as concordant or discordant.

### Aging rate indicators (ARIs)

Candidate ARIs and their directions in slow-aging mice were taken from Austad, Kaeberlein and Miller (2026). Transcript-level ARIs (*DCX, BDNF, FNDC5, GPLD1, UCP1*) were tested as individual genes using the pooled limma model. Pathway-level ARIs were scored as transcriptional activity: mTORC1 using the MSigDB Hallmark MTORC1_SIGNALING set, and MEK1/ERK using MPAS, as described above. ARIs not assayable from a neuronal transcriptome were excluded.

To assess compounds with few individually significant ARI changes, each indicator was scored per sample (log2 CPM for transcript-level; mean z-score for pathway level), and its effect relative to vehicle was estimated with lm(value ∼ treatment + cell line), reported with 95% confidence intervals and nominal p-values without correction for multiple testing. For composite scores, each indicator was centered on the vehicle mean, scaled by its residual SD and sign-aligned; the composite for each sample was the mean across indicators and was tested with the same model. Directional consistency was summarized as the number of indicators shifted in the slow-aging direction. A composite was described as a coordinated shift when its 95% confidence interval excluded zero, and as no detectable response when the entire interval lay within ±0.5 residual SDs.

### Transcriptomic clocks

To remove non-expressed genes from the bulk RNA-seq data set, only those with at least 10 reads in at least 20% of samples were retained. The filtered data were then subjected to Relative Log Expression normalization, log transformation, and YuGene normalization. Missing expression values for clock genes absent from the data set were imputed using their respective precomputed average values. Resulting normalized expression profiles were centered to the median profile of vehicle-treated samples. Transcriptomic age (tAge) and mortality tAge for each sample were estimated using composite elastic net-based multi-species, multi-tissue transcriptomic clocks of chronological age and expected mortality (Tyshkovskiy *et al*., 2026).

### Brain aging signatures

Lu et al. (2004) frontal cortex aging genes were taken from MSigDB C2 (LU_AGING_BRAIN_UP, LU_AGING_BRAIN_DN). Catching et al. (2026) excitatory neuron age-associated genes were taken from Table S3, restricted to the ExN cell type. The PsychAD excitatory-neuron signature was derived here (**Table S2**). Nuclei annotated as glutamatergic neurons from non-diseased donors in the PsychAD lifespan atlas (CELLxGENE dataset 8c3ae5ad-d4c1-4cbe-a8b8-7c5f84bb0664) were aggregated to per-donor pseudobulk counts. Donors aged > 20 years with at least 50 excitatory neurons were retained. Age-associated genes were identified with limma-voom using the model ∼ age_decades + sex + log(n_cells), and the signature comprises genes at FDR < 0.05, capped at 250 per direction. Compound effects on each signature were tested by fgsea on the same moderated t-statistics, with each signature arm treated as a separate gene set and correction applied across compounds within each arm. A combined test weighting age-up genes +1 and age-down genes -1 was run with fry, which accounts for inter-gene correlation.

## Results

### iPSC-derived neurons report compound pharmacology faithfully and reproducibly

iPSC-neurons from one male and one female iPSC line were treated for 48 hours with one of eight compounds or vehicle and profiled by bulk RNA-sequencing in duplicate per line across two independent differentiations (**Figure 1a**). PC1 explained 78.8% of the variance and separated the two donor lines (**Figure S1a**). All subsequent comparisons therefore model treatment with cell-line as a blocking factor. With the line effect removed, trametinib separated clearly from all other conditions, while the remaining compounds overlapped with the vehicle control **(Figure 1b**).

**Figure 1.**
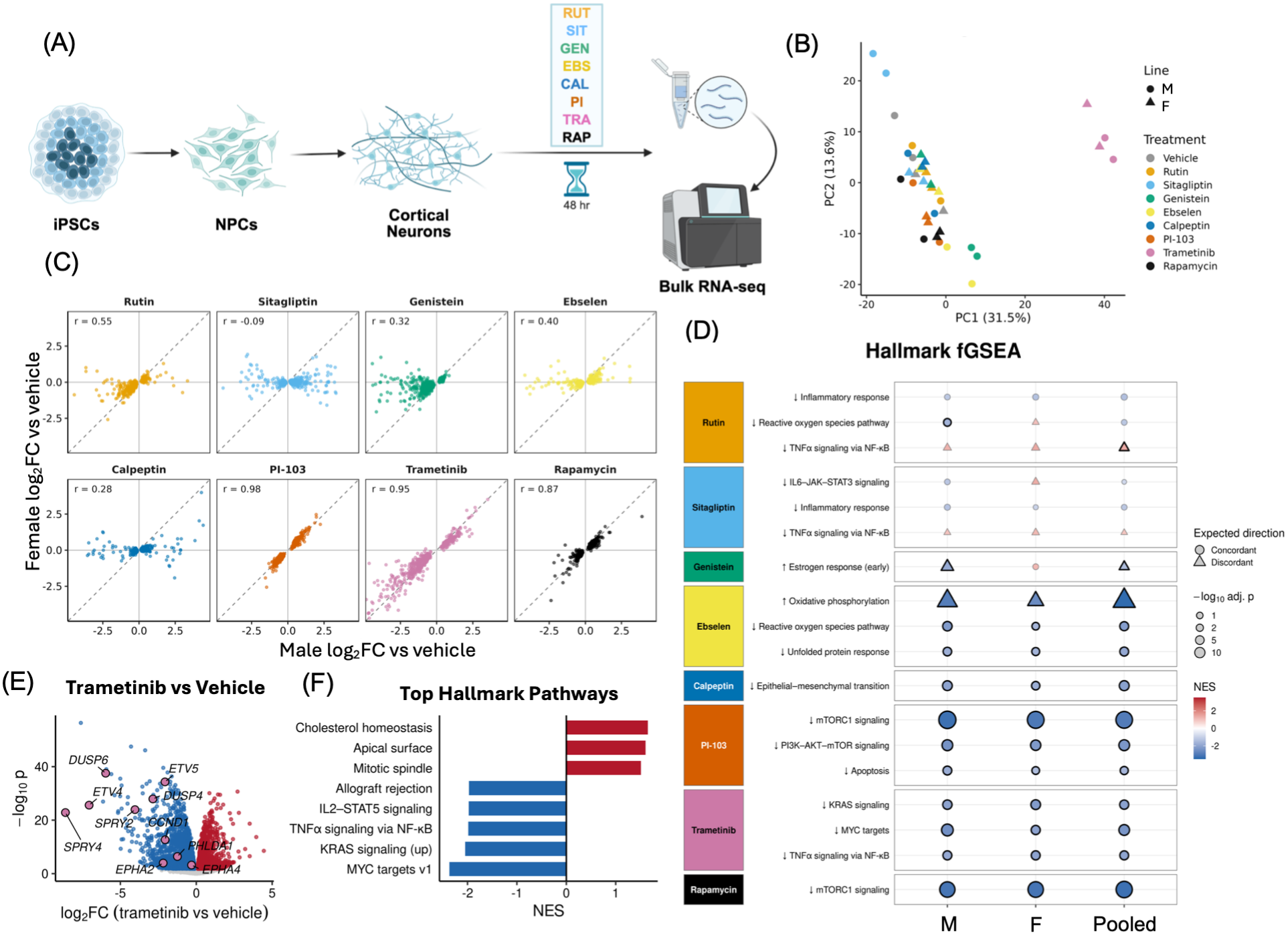
iPSC-derived cortical neurons report geroprotector pharmacology faithfully and reproducibly. (a) Screen design. iPSCs from one male and one female donor were differentiated through neural progenitors to NGN2-induced cortical neurons, matured for 14 days, treated for 48 h with one of eight candidate geroprotectors or vehicle, and profiled by bulk RNA-seq. One replicate from each of two independent differentiations per line (n = 2 per line; n = 4 independent cultures per compound). (b) Principal component analysis after removal of the genetic background effect, using the 500 most variable genes. Color indicates treatment, shape indicates donor. (c) Cross-line reproducibility. For each compound, the 500 most significant genes in one line are plotted as log2 fold change versus vehicle in that line against the same genes measured in the other line. Spearman correlation inset; dashed line is the identity. (d) Hallmark gene set enrichment against the pathway each compound was predicted to modulate (Table 1). Point fill is normalized enrichment score, point size is -log10 Benjamini-Hochberg adjusted p, corrected across the full 50-set Hallmark collection within each contrast. Circles indicate enrichment concordant with the prespecified direction, triangles discordant. Black outline adjusted p < 0.05. (e) Volcano plot of trametinib versus vehicle, pooled model. The ten MAPK Pathway Activity Score genes are highlighted and labelled; all ten are suppressed. Red and blue indicate FDR < 0.05 up and down respectively. (f) Hallmark pathways most enriched by trametinib, ranked by normalized enrichment score, at adjusted p < 0.05.

Compounds differed markedly in the extent of transcriptional response. At FDR < 0.05, trametinib and PI-103 altered 6,054 and 5,438 genes respectively, and rapamycin 2,216. Rutin, genistein and ebselen altered between 635 and 1,602 genes (**Figure S1b, Figure S1d**). Sitagliptin resulted in no individual genes that met the prespecified FDR threshold at the concentrations and duration tested. Effects also differed in reproducibility across donor lines: correlations were 0.98 for PI-103, 0.95 for trametinib and 0.87 for rapamycin (**Figure 1c**).

Hallmark gene set enrichment was tested against the pathway each compound was predicted to modulate (**Table 1**). PI-103 suppressed mTORC1 and PI3K-AKT-mTOR signaling, rapamycin suppressed mTORC1 signaling, and trametinib suppressed KRAS signaling, MYC targets and TNFα signaling via NF-κB (**Figure 1d**). PI-103 and rapamycin each reduced an mTORC1 signature score (adjusted p < 0.001), and trametinib reduced a MEK-ERK transcriptional activity score (MPAS) by approximately 2.5 standard deviations (adjusted p < 0.001), the largest single effect in the screen (**Figure 2b**).

**Figure 2.**
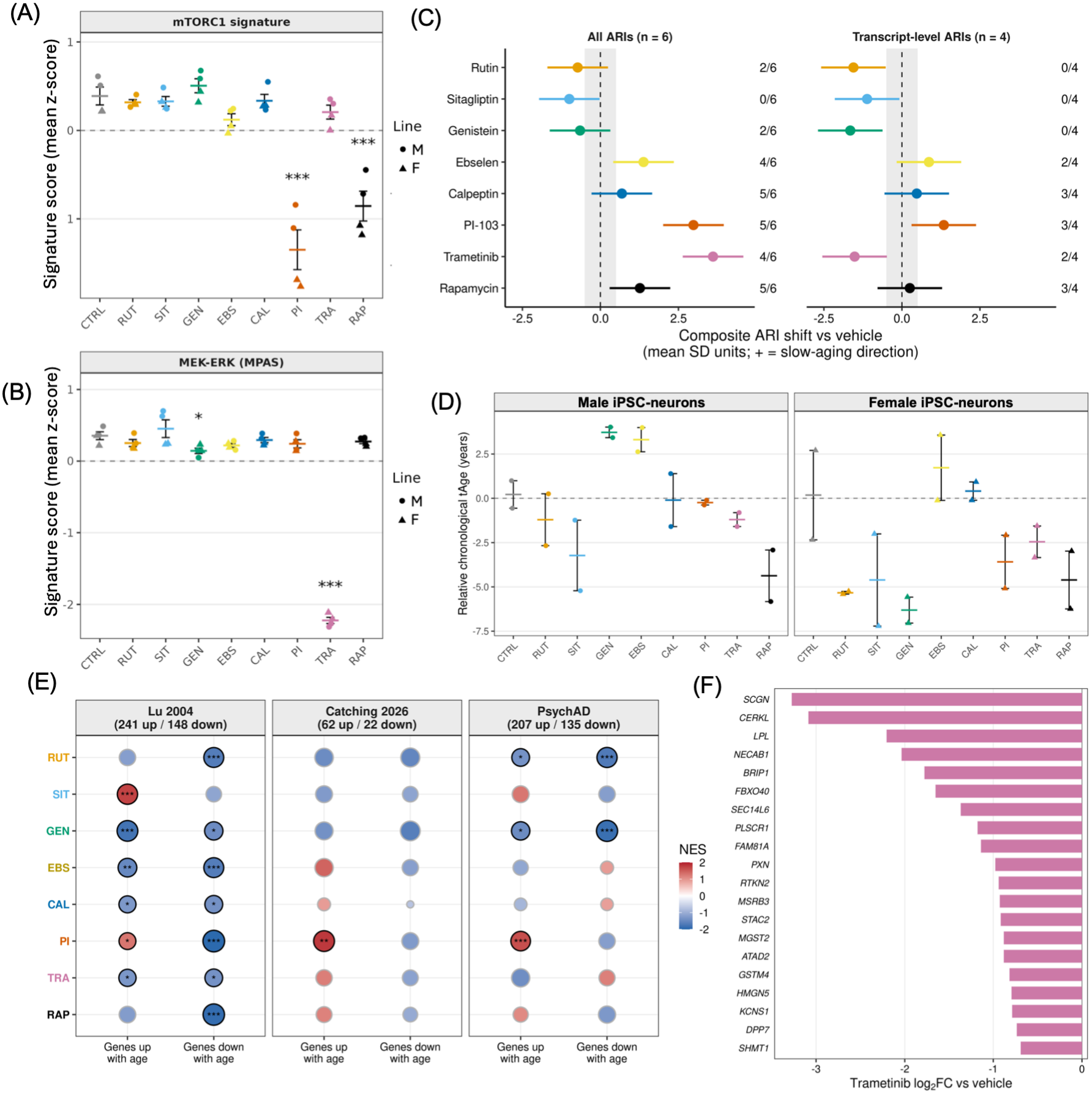
Aging rate indicator, transcriptomic age, and brain aging signature responses across the compound panel. (a,b) Transcriptional activity scores for the two pathway-level aging rate indicators, mTORC1 (a) and MEK1/ERK (b), expressed as the mean z-score across genes. Crossbar is the group mean, error bars are SEM, points are individual samples shaped by donor. Reduced activity is the slow-aging direction for both indicators. Asterisks are from lm(score ∼ treatment + donor), Benjamini-Hochberg corrected. (c) Composite aging rate indicator shift for each compound versus vehicle, calculated across all six ARIs (left) and across the four transcript-level ARIs (right). Each indicator was expressed in residual standard deviation units and sign aligned so that positive values indicate the slow-aging direction. Points are estimated from lm(composite ∼ treatment + donor) and bars are 95% confidence intervals, without correction for multiple testing. (d) Relative chronological transcriptomic age shown separately for the male and female lines. Crossbar is the mean of two independent differentiations, error bars are SEM. (e) Enrichment of each compound against three human brain aging references: bulk frontal cortex (Lu et al., 2004), excitatory neurons from a single-nucleus multiome cohort (Catching et al., 2026), and an excitatory-neuron signature derived here from the PsychAD lifespan atlas. Gene counts per reference are given in the panel headers. Point fill is normalized enrichment score and size is effect magnitude; black outline and asterisks indicate adjusted p < 0.05, corrected across compounds within each signature arm. Blue on the age-up arm and red on the age-down arm correspond to a younger-like shift. (f) The twenty genes most strongly suppressed by trametinib among the leading edge of the PsychAD age-up gene set, plotted as pooled log2 fold change versus vehicle.

At gene level, trametinib suppressed all ten MPAS genes, with *SPRY4, ETV4* and *DUSP6* among the largest effects in the dataset (**Figure 1e, 1f**). The pathways that lifespan studies have identified as longevity-associated are therefore intact in fetal-like human neurons and can be engaged pharmacologically. Three predicted effects were not observed (**Figure 1d**). Ebselen shifted oxidative phosphorylation opposite to prediction in both lines, genistein did likewise for the early estrogen response, and sitagliptin increased rather than decreased IL6-JAK-STAT3 and TNFα/NF-κB signaling.

### Three compounds shift pathway-level aging rate indicators in the slow-aging direction

The screen was scored against twelve candidate ARIs (Austad, Kaeberlein and Miller, 2026), restricted to those that neurons express. Based on these criteria, six ARIs were measurable. *DCX* and *BDNF* are hippocampal indicators in mice. *FNDC5* is a muscle indicator whose cleavage product irisin is the circulating form. *GPLD1* is a hepatic indicator also detectable in plasma. Two further indicators are pathway-level: mTORC1 activity and MEK1/ERK activity, scored here from transcriptional output (**Figure 2a, 2b; Figure S2a**). The pathway-level indicators moved strongly and in the slow-aging direction. PI-103 and rapamycin each reduced the mTORC1 signature (adjusted p < 0.001; **Figure 2a**). Trametinib reduced the MEK-ERK activity score by approximately 2.5 standard deviations (adjusted p < 0.001; **Figure 2b**). Three of eight compounds therefore induced the pathway state that distinguishes slow-aging mice from controls.

The two hippocampal indicators, *DCX* and *BDNF*, extend this result in iPSC-neurons for one compound and qualify it for another (**Figure S2a**). PI-103 increased both *DCX* and *BDNF*, concordant with the slow-aging direction on both, giving it three concordant ARIs in total. Trametinib increased *DCX* but reduced *BDNF* (each adjusted p < 0.001, **Figure S2a**), concordant on one and discordant on the other. Ebselen increased *DCX*, while rutin and genistein reduced *FNDC5*, discordant with the slow-aging direction. Aging rate indicators are therefore expressed at the transcript level, measurable, and pharmacologically responsive in iPSC-neurons that carry no chronological age, and a subset of compounds moves them in the direction that defines a slow-aging state.

Because several compounds produced few individually significant ARI changes, we next asked whether a combined analysis could distinguish a weak but coordinated response from no clear response. Across all six ARIs, four compounds produced composite shifts in the slow-aging direction with confidence intervals excluding zero: trametinib (3.61 SD, 95% CI 2.64 to 4.58; 4 of 6 indicators in the slow aging direction), PI-103 (2.98 SD, 2.01 to 3.95; 5 of 6), ebselen (1.38 SD, 0.41 to 2.35, 4 of 6) and rapamycin (1.27 SD, 0.29 to 2.24; 5 of 6) (**Figure 2c, Figure S2c)**. Calpeptin moved five of six indicators in the slow-aging direction, but its composite interval included zero, consistent with a weak yet coordinated response.

Restricting the composite to the four transcript-level ARIs revealed that PI-103 remained in the slow-aging direction whereas the trametinib composite reversed, driven largely by its reduction in BDNF (**Figure 2c)**. The combined analysis therefore distinguished several response types among compounds with few individually significant changes: weak but coordinated shifts in the slow-aging direction (ebselen), coordinated shifts in the opposite direction (sitagliptin across all ARIs), and responses monopolized by a single pathway indicator (rapamycin).

### Chronological transcriptomic age tracks pathway engagement

We next asked whether transcriptomic clocks return interpretable values in neurons in which reprogramming has reset much of the donor-associated aging signature. Relative chronological tAge and mortality transcriptomic age were estimated using multi-species clocks (Tyshkovskiy *et al*., 2026), with vehicle samples as the reference group. Seven of eight compounds reduced chronological tAge relative to vehicle. Rapamycin produced the largest reduction (-4.7 years), followed by sitagliptin (-4.1) and rutin (-3.5), with PI-103 (-2.1) and trametinib (-2.0) intermediate. Ebselen was the only compound to increase chronological tAge in both cell lines (+2.3) (**Figure 2d, Figure S2b**). All three compounds that shifted pathway-level ARIs in the slow-aging direction (PI-103, trametinib, and rapamycin) reduced chronological tAge. This indicates that a relative clock retains discriminative information when applied to an iPSC culture system.

### Brain aging references differ in composition, and compound effects vary across them

Compounds were then scored against three human brain aging references: bulk frontal cortex (Lu *et al*., 2004), excitatory neurons from a single-nucleus multiome cohort (Catching *et al*., 2026), and an excitatory-neuron signature derived from the PsychAD lifespan atlas (Yang *et al*., 2026). Of the 85 genes shared between the two neuron references, all 85 change in the same direction with age (binomial p = 5x10^-26^, rho = 0.58; **Figure S2e**). The bulk reference agrees with neither, sharing almost no genes with either neuron set and matching direction on only 61% of shared genes (**Figure S2g**).

Compound effects also varied across references (**Figure 2e**). PI-103 was the only compound called older-like by both neuron-derived references, significantly enriching age-up genes in each. Trametinib showed a non-significant trend toward suppression of age-up genes on the PsychAD reference, consistent with a younger-like shift, driven by a leading edge of 61 of the 207 expressed age-up genes, with *SCGN, CERKL* and *LPL* among the strongest (**Figure 2f**). No compound produced a consistent younger-like shift across references, and none realigned the transcriptome against neuronal aging at genome scale, with correlations between each compound’s gene-level effects and the PsychAD age effects ranging from rho = −0.02 to 0.11.

## Discussion

We set out to explore whether a human iPSC-system can be used as a model to screen for geroprotectors. Longevity-associated pathways are intact in iPSC-derived neurons and respond to pharmacological manipulation with effects large enough to detect and reproducible enough to generalize across donors and differentiations. Six of twelve candidate aging rate indicators are measurable in these cells, and three compounds (PI-103, rapamycin, and trametinib) moved a pathway ARI in the direction that distinguishes slow-aging mice from controls. This establishes expected pharmacology and a geroprotection-associated response, but pathway level engagement alone does not demonstrate protection against aging.

These results were echoed by the transcriptomic clock interpretation of these data as well, where seven of eight compounds reduced chronological tAge. Rapamycin ranked first, the transcriptionally inert compound sat at zero, and the one compound with a discordant pathway profile was the only one to move in the opposite direction. All three ARI-positive compounds reduced chronological tAge. A relative clock, referenced against vehicle within the same experiment, therefore appears to retain discriminative information in a rejuvenated system. In contrast, human brain aging signatures, which are derived from tissue that has accumulated age-associated molecular and compositional changes, ranked compounds differently depending on the reference applied. The combined ARI analysis added a further example of disagreement; ebselen produced a coordinated shift across all ARIs in the slow aging direction (**Figure 2c**) but was the only compound to increase chronological tAge (**Figure 2d**). Rather than indicating that one transcriptional readout is correct, these divergences suggest that each captures a different aspect of drug response.

The throughput argument is straightforward. While a mouse lifespan study takes three to four years, the present screen took 48 hours of compound exposure and returned human tissue-specific pathway-level, gene-level and indicator-level readouts. The comparison is not intended to suggest that culture readouts replace lifespan studies but instead aims to emphasize the utility of these systems for larger screens to prioritize compounds for subsequent *in vivo* lifespan studies.

A practical implication of this work is that bulk RNA-seq may serve as a scalable proxy for several ARIs that are conventionally measured by protein or signaling assays. If transcript level ARI scores track their protein-level counterparts, RNA-seq could allow ARI panels to be applied across many compounds, doses, and donors at a fraction of the cost. This correspondence has not yet been established, and for the present model, paired RNA-seq and protein or phosphor-protein measurements in the same samples will be needed to validate transcript-level ARIs before they are used as a proxy.

A further advantage of iPSC-based systems for evaluating candidate geroprotectors is the ability to sample human genetic diversity across many donors. In the present screen, we see evidence of this at play as reflected in the donor-discordant tAge shift with genistein (**Figure 2d**). This is particularly relevant to sex as a biological variable. Most of the compounds identified by the ITP preferentially extend lifespan in male UM-HET3 mice (Jiang *et al*., 2025), and the basis of this sexual divergence remains poorly understood. Because iPSC lines can be derived from large, sex-balanced donor cohorts, they offer a scalable human model for dissecting sex-dependent responses to geroprotective compounds across tissues of all lineages. In the present study, we compared a single male and a single female donor, which does not allow sex-specific effects to be distinguished from inter-individual genetic variation. Additional studies incorporating multiple donors of each sex will be needed to determine whether geroprotector efficacy diverges by sex in human neurons and across genetic backgrounds.

Future studies could build on this screen in several ways. Directly converted neurons retain donor age signatures (Mertens *et al*., 2018), so running the same panel in aged neurons would allow ARI and biomarker-type readouts to be compared in cells that do carry age-associated features. Notably, in age- and disease-retaining iSpheroids, trametinib and PI-103 produced the largest transcriptional responses among 12 compounds screened by bulk RNA-sequencing (Alsolami *et al*., 2026). These results are strikingly similar to what we observed in our 2D rejuvenated iPSC-neuron screen. This suggests that, at the transcriptional level, geroprotector screening may elicit similar top-contenders in aged vs. rejuvenated systems of the same tissue. Applying the same compounds to iPSC-derived hepatocytes, myocytes and adipocytes would test tissue-specific geroprotection directly, using indicators that neurons cannot report, such as *UCP1*. Future screens would also benefit from negative controls such as resveratrol and simvastatin, which have been identified by the ITP as non-lifespan extending (Miller *et al*., 2011).

Trametinib produced the largest transcriptional response in the panel, and the largest single pathway ARI effect. However, the trametinib composite across transcript-level ARIs moved away from the slow-aging direction, so its nomination in human neurons will require confirmation with functional readouts. In summary, geroprotector screening has been constrained by an assay that takes years and reports a single organism-level endpoint. Human iPSC-derived cells remove that constraint, and the transcriptional readouts now available to the aging field make the resulting data interpretable.

Several limitations apply. One male and one female line do not permit sex to be separated from donor, and several compounds produced effects of opposite sign and of various magnitudes in the two lines. This is a constraint of the present screen rather than of the platform, since donor number is precisely the variable iPSC systems can scale. No functional or survival endpoints were measured. The combined ARI analysis was exploratory: effects were evaluated without correction for multiple testing, composite scores depend on how indicators are weighted, and all indicators were derived from bulk RNA-seq data instead of protein. The pathway activity scores measure transcriptional output of mTORC1 and MEK-ERK signaling rather than kinase activity, which would require phospho-protein measurement to establish. Finally, compounds with weak or absent signals cannot yet be confidently deprioritized. Sitagliptin and calpeptin produced few or no DEGs, and several compounds showed composite ARI shifts whose confidence intervals included zero. These results may represent false negatives arising from the single concentration, acute exposure, or the neuronal context rather than an absence of activity. Dose-response experiments across multiple lineages derived from the same donor, together with protein level and functional assays, would help distinguish inactive compounds from those tested outside their effective range in the present study.

## Data code & availability

Data are available on GEO under repository number GSE354186. The analysis code used in this study is available on GitHub at https://github.com/elisemac-crem/GeroScreen_2026.

## Acknowledgments

We thank the members of the Longevity Consortium for their assistance in selecting compounds to prioritize in the present targeted screen.

## Author Contributions

Conceptualization: EMM, GJM

Methodology: EMM

Investigation: EMM, NL, NJ Skvir, CVM, SFC

Visualization: EMM, CVM

Funding Acquisition: GJM, PS

Writing – original draft: EMM

Writing – review and editing: EMM, TG, SFC, NL, CVM, NJ Schork, GJM

## Figure Legends

**Supplementary Figure 1. Transcriptional response magnitude and structure across the screen**. (a) Principal component analysis of all samples before correction, using the 500 most variable genes. PC1 separates the two donors and accounts for 78.8% of variance, which is why all comparisons in the main text block on cell line. (b) Differentially expressed genes per compound versus vehicle at Benjamini-Hochberg adjusted p < 0.05, split by direction. (c) Sample-to-sample Pearson correlation after removal of the genetic background effect, over the 500 most variable genes, hierarchically clustered. BU6 is the male line, BU8 is the female line. The colored strip indicates treatment. (d) Volcano plots for each compound versus vehicle, pooled model, with shared axes so effect magnitudes are directly comparable. Colored points are FDR < 0.05; the three most significant genes per direction are labelled. Counts of up- and down-regulated genes are inset.

**Supplementary Figure 2. Aging readouts, quality control and reference validation**. (a) Transcript-level aging rate indicators (DCX, BDNF, GPLD1, FNDC5) shown as log2 fold change versus vehicle for each compound, with individual samples shaped by donor. UCP1 is omitted because it was not detectably expressed. (b) Relative chronological and mortality transcriptomic age for each compound, pooled across donors and adjusted for cell line. Crossbar is the group mean, error bars are SEM. Asterisks are from lm(tAge ∼ treatment + donor), Benjamini-Hochberg corrected across compounds. (c) Effect of each compound on each ARI versus vehicle, expressed in residual standard deviation units and sign-aligned so that positive values indicate the slow-aging direction. Filled points indicate nominal p < 0.05, without correction for multiple testing. (d) Absolute chronological tAge shift plotted against the number of differentially expressed genes for each compound, on a log scale with zero plotted at the axis origin. Sitagliptin produces the second-largest clock shift with no differentially expressed genes. (e) Validation of the PsychAD signature against the independent Catching et al. (2026) excitatory-neuron age genes. Each point is a gene shared between the two references, plotted by its age effect in each. Shaded quadrants indicate directional agreement. (f) Multidimensional scaling of per-donor excitatory-neuron pseudobulk from the PsychAD lifespan atlas, adult donors only, colored by donor age and sized by the number of excitatory neurons contributing to each donor. (g) Genes shared between each pair of signature arms across the three brain aging references. Diagonal entries in bold give the size of each set; fill indicates the percentage of the row set shared with the column set.

