## Supplemental Table 1 and Supplemental Figures for "Comparative transcriptomic validation of candidate geroprotectors in human iPSC-derived neurons"

#### Supplementary Table 1

**Table S1. Compounds, concentrations, and sources used in the screen.** Stock and final concentrations, supplier, and catalogue number for each compound. NGN2-induced cortical neurons were matured for 14 days before treatment and treated for 48 hours.

Stocks were prepared in DMSO and diluted to a 1 mM (or 100 μM) intermediate before addition to medium; vehicle wells received an equivalent volume of DMSO alone.

| **Compound** | **Abbreviation** | **Stock concentration** | **Final concentration** | **Supplier** | **Catalogue no.** |
|---|---|---|---|---|---|
| Rutin | RUT | 10 mM | 5 μM | Selleck Chemicals | S2350 |
| Sitagliptin | SIT | 10 mM | 1 μM | Selleck Chemicals | S5079 |
| Genistein | GEN | 10 mM | 5 μM | Selleck Chemicals | S1342 |
| Ebselen | EBS | 10 mM | 1 μM | Selleck Chemicals | S6676 |
| Calpeptin | CAL | 10 mM | 1 μM | Selleck Chemicals | S7396 |
| PI-103 | PI | 5 mM | 500 nM | Selleck Chemicals | S1038 |
| Trametinib | TRA | 5 mM | 50 nM | Selleck Chemicals | S2673 |
| Rapamycin | RAP | 1 mM | 50 nM | Sigma-Aldrich | R8781 |

**Supplementary Table 2 (*TableS2.EMM.pdf*)**

**Table S2. Genes associated with age in excitatory neurons from non-diseased adult donors in the PsychAD lifespan atlas (Yang et al., 2026; CELLxGENE dataset 8c3ae5ad-d4c1-4cbe-a8b8-7c5f84bb0664).** Per-donor pseudobulk counts were generated from nuclei annotated as glutamatergic neurons, retaining donors aged 20 years or older with at least 50 excitatory neurons. Age associations were tested with limma-voom using the model ~ age (per decade) + sex + log(number of cells). The signature comprises genes at FDR < 0.05, capped at 250 per direction (250 up with age, 250 down with age).

### Supplementary Figure 1

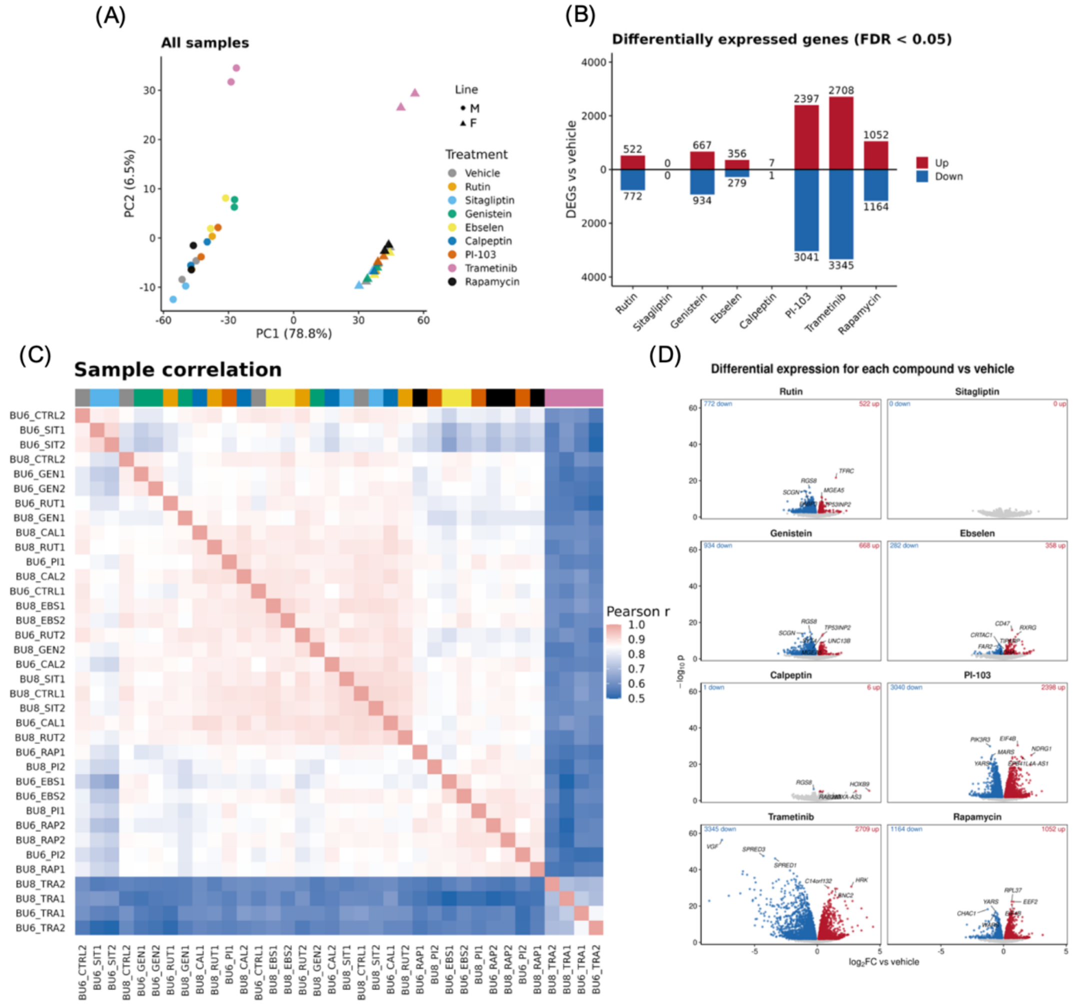

### Supplementary Figure 2

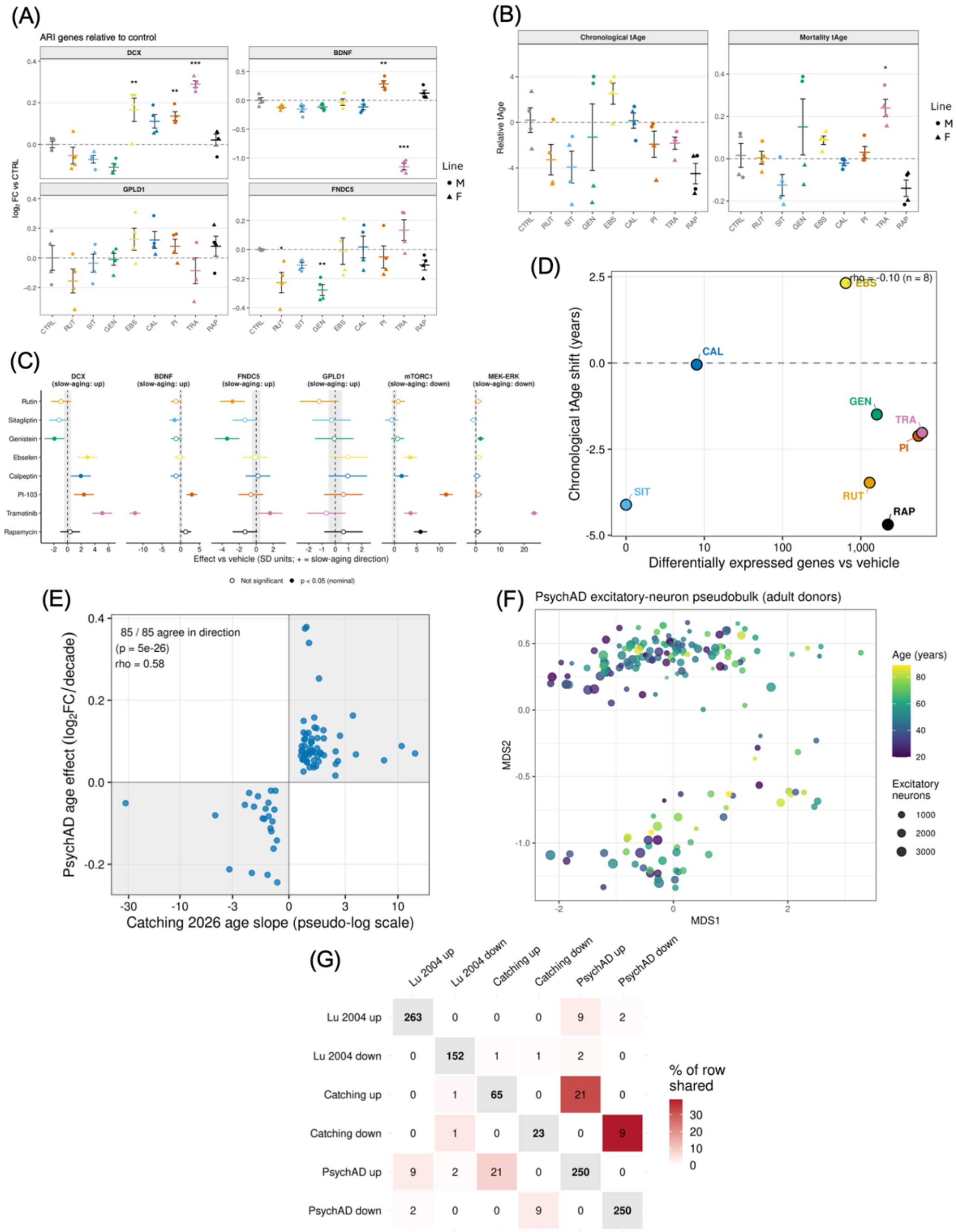


| | Lu 2004 up | Lu 2004 down | Catching up | Catching down | PsychAD up | PsychAD down |
|---|---|---|---|---|---|---|
| Lu 2004 up | 263 | 0 | 0 | 0 | 9 | 2 |
| Lu 2004 down | 0 | 152 | 1 | 1 | 2 | 0 |
| Catching up | 0 | 1 | 65 | 0 | 21 | 0 |
| Catching down | 0 | 1 | 0 | 23 | 0 | 9 |
| PsychAD up | 9 | 2 | 21 | 0 | 250 | 0 |
| PsychAD down | 2 | 0 | 0 | 9 | 0 | 250 |
