## Supplemental Table 2 for "Comparative transcriptomic validation of candidate geroprotectors in human iPSC-derived neurons"

### Supplementary Table S2. PsychAD excitatory-neuron aging signature.

Genes associated with age in excitatory neurons from non-diseased adult donors in the PsychAD lifespan atlas (Yang et al., 2026; CELLxGENE dataset 8c3ae5ad-d4c1-4cbe-a8b8-7c5f84bb0664). Per-donor pseudobulk counts were generated from nuclei annotated as glutamatergic neurons, retaining donors aged 20 years or older with at least 50 excitatory neurons. Age associations were tested with limma-voom using the model ~ age (per decade) + sex + log(number of cells). The signature comprises genes at FDR < 0.05, capped at 250 per direction (250 up with age, 250 down with age). Log2 fold changes are expressed per decade of age. Up-regulated genes are listed first; within each direction, genes are ordered by FDR.

| Gene | Direction with age | log2 fold change per decade | Average log2 CPM | Moderated t | P value | FDR |
|---|---|---|---|---|---|---|
| *BMP4* | Up | 0.498 | –0.68 | 14.25 | $2.56 \times 10^{-31}$ | $7.26 \times 10^{-27}$ |
| *RHBDL3* | Up | 0.340 | 5.56 | 13.48 | $4.72 \times 10^{-29}$ | $5.80 \times 10^{-25}$ |
| *DOCK11* | Up | 0.171 | 2.99 | 12.47 | $4.11 \times 10^{-26}$ | $1.95 \times 10^{-22}$ |
| *FKBP5* | Up | 0.379 | 5.03 | 12.35 | $9.50 \times 10^{-26}$ | $3.77 \times 10^{-22}$ |
| *RPS10P7* | Up | 0.226 | 2.04 | 11.62 | $1.21 \times 10^{-23}$ | $3.43 \times 10^{-20}$ |
| *USP54* | Up | 0.135 | 5.57 | 11.32 | $9.23 \times 10^{-23}$ | $2.39 \times 10^{-19}$ |
| *ROS1* | Up | 0.375 | 1.65 | 10.92 | $1.28 \times 10^{-21}$ | $2.60 \times 10^{-18}$ |
| *SPOCK3* | Up | 0.253 | 6.37 | 10.21 | $1.41 \times 10^{-19}$ | $1.90 \times 10^{-16}$ |
| *PKD1L1* | Up | 0.224 | 4.65 | 10.13 | $2.25 \times 10^{-19}$ | $2.78 \times 10^{-16}$ |
| *GOLGA2P5* | Up | 0.204 | 1.58 | 10.09 | $2.92 \times 10^{-19}$ | $3.31 \times 10^{-16}$ |
| *CLGN* | Up | 0.153 | 2.57 | 9.90 | $9.98 \times 10^{-19}$ | $1.01 \times 10^{-15}$ |
| *HEBP2* | Up | 0.273 | 0.57 | 9.59 | $7.65 \times 10^{-18}$ | $7.01 \times 10^{-15}$ |
| *C2CD2* | Up | 0.152 | 4.64 | 9.41 | $2.34 \times 10^{-17}$ | $1.96 \times 10^{-14}$ |
| *STAT4* | Up | 0.158 | 6.72 | 9.40 | $2.47 \times 10^{-17}$ | $2.00 \times 10^{-14}$ |
| *FUT10* | Up | 0.134 | 3.91 | 9.36 | $3.21 \times 10^{-17}$ | $2.54 \times 10^{-14}$ |
| *LAMA3* | Up | 0.178 | 2.78 | 9.33 | $3.93 \times 10^{-17}$ | $3.02 \times 10^{-14}$ |
| *SLC26A11* | Up | 0.114 | 4.11 | 9.33 | $4.05 \times 10^{-17}$ | $3.03 \times 10^{-14}$ |
| *CCND3* | Up | 0.194 | 5.47 | 9.08 | $1.89 \times 10^{-16}$ | $1.22 \times 10^{-13}$ |
| *SERPING1* | Up | 0.133 | 2.53 | 9.07 | $2.02 \times 10^{-16}$ | $1.27 \times 10^{-13}$ |
| *DMKN* | Up | 0.133 | 3.23 | 9.03 | $2.55 \times 10^{-16}$ | $1.48 \times 10^{-13}$ |
| *SMCO4* | Up | 0.267 | 2.84 | 9.00 | $3.14 \times 10^{-16}$ | $1.78 \times 10^{-13}$ |
| *CHMP1B2P* | Up | 0.267 | 1.55 | 8.96 | $3.99 \times 10^{-16}$ | $2.14 \times 10^{-13}$ |
| *KDSR* | Up | 0.084 | 4.50 | 8.87 | $7.35 \times 10^{-16}$ | $3.67 \times 10^{-13}$ |
| ENSG00000253642 | Up | 0.176 | 3.25 | 8.85 | $8.21 \times 10^{-16}$ | $4.02 \times 10^{-13}$ |
| *UPP1* | Up | 0.089 | 4.60 | 8.84 | $8.64 \times 10^{-16}$ | $4.16 \times 10^{-13}$ |
| *RCL1* | Up | 0.136 | 4.89 | 8.81 | $1.03 \times 10^{-15}$ | $4.89 \times 10^{-13}$ |
| *CAPN2* | Up | 0.093 | 4.10 | 8.74 | $1.66 \times 10^{-15}$ | $7.73 \times 10^{-13}$ |
| *MAP3K20* | Up | 0.165 | 3.81 | 8.69 | $2.16 \times 10^{-15}$ | $9.89 \times 10^{-13}$ |
| *ANK1* | Up | 0.208 | 3.53 | 8.68 | $2.38 \times 10^{-15}$ | $1.07 \times 10^{-12}$ |
| *ENDOD1* | Up | 0.083 | 4.28 | 8.63 | $3.24 \times 10^{-15}$ | $1.42 \times 10^{-12}$ |
| *WFS1* | Up | 0.151 | 2.18 | 8.53 | $5.79 \times 10^{-15}$ | $2.46 \times 10^{-12}$ |
| *CALCOCO2* | Up | 0.103 | 5.33 | 8.52 | $6.16 \times 10^{-15}$ | $2.57 \times 10^{-12}$ |
| *SERPINB6* | Up | 0.130 | 3.75 | 8.51 | $6.76 \times 10^{-15}$ | $2.74 \times 10^{-12}$ |
| *HELLS* | Up | 0.112 | 3.44 | 8.44 | $1.03 \times 10^{-14}$ | $4.00 \times 10^{-12}$ |
| *EPRS1* | Up | 0.076 | 5.52 | 8.42 | $1.13 \times 10^{-14}$ | $4.29 \times 10^{-12}$ |
| *GPC5* | Up | 0.166 | 7.51 | 8.41 | $1.24 \times 10^{-14}$ | $4.57 \times 10^{-12}$ |
| ENSG00000242593 | Up | 0.156 | 7.56 | 8.36 | $1.69 \times 10^{-14}$ | $6.08 \times 10^{-12}$ |
| ENSG00000257489 | Up | 0.180 | 0.73 | 8.34 | $1.86 \times 10^{-14}$ | $6.44 \times 10^{-12}$ |
| *YPEL4* | Up | 0.103 | 3.57 | 8.33 | $1.96 \times 10^{-14}$ | $6.72 \times 10^{-12}$ |
| *FAM81A* | Up | 0.107 | 6.83 | 8.31 | $2.31 \times 10^{-14}$ | $7.81 \times 10^{-12}$ |
| *INPP4B* | Up | 0.164 | 8.26 | 8.30 | $2.47 \times 10^{-14}$ | $8.26 \times 10^{-12}$ |
| ENSG00000259403 | Up | 0.206 | 3.39 | 8.29 | $2.57 \times 10^{-14}$ | $8.48 \times 10^{-12}$ |
| ENSG00000287037 | Up | 0.236 | –0.43 | 8.22 | $3.85 \times 10^{-14}$ | $1.23 \times 10^{-11}$ |
| *NT5M* | Up | 0.094 | 3.73 | 8.20 | $4.46 \times 10^{-14}$ | $1.38 \times 10^{-11}$ |
| *BRIP1* | Up | 0.136 | 3.37 | 8.19 | $4.61 \times 10^{-14}$ | $1.41 \times 10^{-11}$ |
| *DZIP1L* | Up | 0.144 | 2.61 | 8.13 | $6.65 \times 10^{-14}$ | $1.97 \times 10^{-11}$ |
| *SLC19A3* | Up | 0.192 | –0.21 | 8.00 | $1.49 \times 10^{-13}$ | $4.15 \times 10^{-11}$ |
| *PTK2B* | Up | 0.134 | 7.25 | 8.00 | $1.51 \times 10^{-13}$ | $4.17 \times 10^{-11}$ |
| *MAGI2-AS3* | Up | 0.114 | 5.99 | 7.99 | $1.55 \times 10^{-13}$ | $4.23 \times 10^{-11}$ |

| Gene | Direction with age | log2 fold change per decade | Average log2 CPM | Moderated t | P value | FDR |
|---|---|---|---|---|---|---|
| *SORBS3* | Up | 0.156 | 2.13 | 7.97 | $1.76 \times 10^{-13}$ | $4.76 \times 10^{-11}$ |
| *BEND7* | Up | 0.163 | 2.82 | 7.97 | $1.79 \times 10^{-13}$ | $4.79 \times 10^{-11}$ |
| *PYGB* | Up | 0.112 | 5.24 | 7.95 | $1.98 \times 10^{-13}$ | $5.22 \times 10^{-11}$ |
| *LINC00173* | Up | 0.152 | 2.03 | 7.94 | $2.14 \times 10^{-13}$ | $5.43 \times 10^{-11}$ |
| *NT5C1A* | Up | 0.128 | 2.73 | 7.93 | $2.22 \times 10^{-13}$ | $5.54 \times 10^{-11}$ |
| *EPB41* | Up | 0.131 | 6.47 | 7.92 | $2.34 \times 10^{-13}$ | $5.77 \times 10^{-11}$ |
| ENSG00000283782 | Up | 0.127 | 4.74 | 7.92 | $2.35 \times 10^{-13}$ | $5.77 \times 10^{-11}$ |
| *EFNA1* | Up | 0.176 | 0.57 | 7.89 | $2.90 \times 10^{-13}$ | $6.99 \times 10^{-11}$ |
| *JAK2* | Up | 0.121 | 6.91 | 7.86 | $3.32 \times 10^{-13}$ | $7.86 \times 10^{-11}$ |
| *LINC01117* | Up | 0.185 | 4.85 | 7.85 | $3.66 \times 10^{-13}$ | $8.59 \times 10^{-11}$ |
| *CIT* | Up | 0.108 | 7.90 | 7.84 | $3.76 \times 10^{-13}$ | $8.71 \times 10^{-11}$ |
| *DKK3* | Up | 0.110 | 6.60 | 7.84 | $3.77 \times 10^{-13}$ | $8.71 \times 10^{-11}$ |
| *CTPS2* | Up | 0.093 | 3.96 | 7.83 | $3.95 \times 10^{-13}$ | $9.05 \times 10^{-11}$ |
| *SYN3* | Up | 0.163 | 9.22 | 7.81 | $4.63 \times 10^{-13}$ | $1.04 \times 10^{-10}$ |
| *N4BP2* | Up | 0.149 | 4.76 | 7.78 | $5.43 \times 10^{-13}$ | $1.20 \times 10^{-10}$ |
| *TMPO* | Up | 0.073 | 4.71 | 7.74 | $6.80 \times 10^{-13}$ | $1.46 \times 10^{-10}$ |
| *LINC02796* | Up | 0.219 | 2.71 | 7.74 | $7.15 \times 10^{-13}$ | $1.52 \times 10^{-10}$ |
| *LYPD5* | Up | 0.096 | 4.10 | 7.66 | $1.11 \times 10^{-12}$ | $2.24 \times 10^{-10}$ |
| *PTPN21* | Up | 0.075 | 4.43 | 7.66 | $1.12 \times 10^{-12}$ | $2.26 \times 10^{-10}$ |
| *GULOP* | Up | 0.263 | –1.13 | 7.65 | $1.16 \times 10^{-12}$ | $2.32 \times 10^{-10}$ |
| *ERICH5* | Up | 0.146 | 0.93 | 7.65 | $1.17 \times 10^{-12}$ | $2.33 \times 10^{-10}$ |
| *TMEM38B* | Up | 0.101 | 3.76 | 7.64 | $1.28 \times 10^{-12}$ | $2.53 \times 10^{-10}$ |
| *WNT3* | Up | 0.148 | 3.91 | 7.63 | $1.34 \times 10^{-12}$ | $2.59 \times 10^{-10}$ |
| *SHMT1* | Up | 0.100 | 2.78 | 7.60 | $1.53 \times 10^{-12}$ | $2.90 \times 10^{-10}$ |
| *NECAB1* | Up | 0.113 | 8.65 | 7.59 | $1.65 \times 10^{-12}$ | $3.10 \times 10^{-10}$ |
| *HAPLN2* | Up | 0.111 | 3.02 | 7.58 | $1.78 \times 10^{-12}$ | $3.34 \times 10^{-10}$ |
| *WIPF3* | Up | 0.172 | 5.84 | 7.56 | $1.98 \times 10^{-12}$ | $3.66 \times 10^{-10}$ |
| *IGDCC3* | Up | 0.148 | 2.92 | 7.55 | $2.12 \times 10^{-12}$ | $3.90 \times 10^{-10}$ |
| *DACH2* | Up | 0.090 | 4.90 | 7.52 | $2.46 \times 10^{-12}$ | $4.46 \times 10^{-10}$ |
| *CREBRF* | Up | 0.074 | 6.18 | 7.51 | $2.60 \times 10^{-12}$ | $4.68 \times 10^{-10}$ |
| *SLFN5* | Up | 0.114 | 2.46 | 7.50 | $2.84 \times 10^{-12}$ | $5.04 \times 10^{-10}$ |
| *MYBPHL* | Up | 0.220 | 1.88 | 7.47 | $3.38 \times 10^{-12}$ | $5.89 \times 10^{-10}$ |
| *SORT1* | Up | 0.120 | 7.10 | 7.42 | $4.61 \times 10^{-12}$ | $7.89 \times 10^{-10}$ |
| *SCGN* | Up | 0.342 | –0.68 | 7.41 | $4.80 \times 10^{-12}$ | $8.11 \times 10^{-10}$ |
| *KAT2B* | Up | 0.110 | 4.92 | 7.40 | $5.03 \times 10^{-12}$ | $8.46 \times 10^{-10}$ |
| *DPP7* | Up | 0.110 | 4.79 | 7.37 | $6.05 \times 10^{-12}$ | $1.01 \times 10^{-9}$ |
| *JHY* | Up | 0.107 | 3.98 | 7.36 | $6.49 \times 10^{-12}$ | $1.05 \times 10^{-9}$ |
| *HUS1* | Up | 0.067 | 3.66 | 7.35 | $6.61 \times 10^{-12}$ | $1.07 \times 10^{-9}$ |
| *GAS1RR* | Up | 0.143 | 3.22 | 7.34 | $7.16 \times 10^{-12}$ | $1.14 \times 10^{-9}$ |
| *IRAK4* | Up | 0.097 | 1.58 | 7.33 | $7.56 \times 10^{-12}$ | $1.19 \times 10^{-9}$ |
| *KDM2B-DT* | Up | 0.129 | 3.83 | 7.32 | $8.07 \times 10^{-12}$ | $1.26 \times 10^{-9}$ |
| *CEACAM19* | Up | 0.101 | 2.84 | 7.30 | $8.88 \times 10^{-12}$ | $1.38 \times 10^{-9}$ |
| *EVA1C* | Up | 0.072 | 5.49 | 7.30 | $9.00 \times 10^{-12}$ | $1.39 \times 10^{-9}$ |
| *TARS3* | Up | 0.059 | 6.32 | 7.30 | $9.10 \times 10^{-12}$ | $1.40 \times 10^{-9}$ |
| *LONRF3* | Up | 0.250 | 2.39 | 7.29 | $9.52 \times 10^{-12}$ | $1.44 \times 10^{-9}$ |
| *CAPN7* | Up | 0.056 | 6.15 | 7.27 | $1.07 \times 10^{-11}$ | $1.58 \times 10^{-9}$ |
| *ITPRID2* | Up | 0.076 | 5.73 | 7.27 | $1.09 \times 10^{-11}$ | $1.60 \times 10^{-9}$ |
| *LINC01695* | Up | 0.197 | 3.03 | 7.25 | $1.17 \times 10^{-11}$ | $1.69 \times 10^{-9}$ |
| *PLSCR1* | Up | 0.123 | 3.49 | 7.25 | $1.17 \times 10^{-11}$ | $1.69 \times 10^{-9}$ |
| *NCKIPSD* | Up | 0.088 | 4.13 | 7.24 | $1.25 \times 10^{-11}$ | $1.78 \times 10^{-9}$ |
| ENSG00000259124 | Up | 0.101 | 3.70 | 7.24 | $1.27 \times 10^{-11}$ | $1.80 \times 10^{-9}$ |
| *IRS1* | Up | 0.120 | 4.20 | 7.23 | $1.30 \times 10^{-11}$ | $1.83 \times 10^{-9}$ |
| *GAB1* | Up | 0.132 | 7.41 | 7.23 | $1.32 \times 10^{-11}$ | $1.84 \times 10^{-9}$ |
| *MGST2* | Up | 0.104 | 4.20 | 7.23 | $1.33 \times 10^{-11}$ | $1.84 \times 10^{-9}$ |
| ENSG00000232855 | Up | 0.170 | 5.64 | 7.21 | $1.49 \times 10^{-11}$ | $2.04 \times 10^{-9}$ |
| *NAAA* | Up | 0.086 | 3.95 | 7.21 | $1.54 \times 10^{-11}$ | $2.09 \times 10^{-9}$ |
| *ELN* | Up | 0.130 | 2.35 | 7.20 | $1.55 \times 10^{-11}$ | $2.10 \times 10^{-9}$ |

| Gene | Direction with age | log2 fold change per decade | Average log2 CPM | Moderated t | P value | FDR |
|---|---|---|---|---|---|---|
| *ALS2CL* | Up | 0.172 | 2.96 | 7.18 | $1.75 \times 10^{-11}$ | $2.35 \times 10^{-9}$ |
| *CXXC4-AS1* | Up | 0.097 | 6.58 | 7.15 | $2.07 \times 10^{-11}$ | $2.72 \times 10^{-9}$ |
| *TNRC6A* | Up | 0.075 | 9.26 | 7.13 | $2.32 \times 10^{-11}$ | $2.99 \times 10^{-9}$ |
| *HAUS5* | Up | 0.113 | 1.94 | 7.13 | $2.38 \times 10^{-11}$ | $3.04 \times 10^{-9}$ |
| *SEMA6A* | Up | 0.134 | 4.66 | 7.12 | $2.45 \times 10^{-11}$ | $3.11 \times 10^{-9}$ |
| *STAC2* | Up | 0.140 | 3.50 | 7.12 | $2.52 \times 10^{-11}$ | $3.18 \times 10^{-9}$ |
| *ISLR* | Up | 0.228 | 0.16 | 7.11 | $2.61 \times 10^{-11}$ | $3.29 \times 10^{-9}$ |
| *LIFR-AS1* | Up | 0.134 | 3.46 | 7.10 | $2.75 \times 10^{-11}$ | $3.43 \times 10^{-9}$ |
| *PMFBP1* | Up | 0.197 | 1.20 | 7.09 | $2.89 \times 10^{-11}$ | $3.59 \times 10^{-9}$ |
| *CENPC* | Up | 0.080 | 6.77 | 7.09 | $2.97 \times 10^{-11}$ | $3.66 \times 10^{-9}$ |
| *KCNS1* | Up | 0.155 | 2.78 | 7.09 | $3.01 \times 10^{-11}$ | $3.71 \times 10^{-9}$ |
| *SAMD14* | Up | 0.149 | 2.69 | 7.08 | $3.12 \times 10^{-11}$ | $3.82 \times 10^{-9}$ |
| *VAMP1* | Up | 0.121 | 3.12 | 7.03 | $4.03 \times 10^{-11}$ | $4.86 \times 10^{-9}$ |
| *SGSH* | Up | 0.110 | 1.42 | 7.03 | $4.22 \times 10^{-11}$ | $5.06 \times 10^{-9}$ |
| *SLC25A13* | Up | 0.069 | 5.01 | 7.01 | $4.62 \times 10^{-11}$ | $5.42 \times 10^{-9}$ |
| *ZBTB20* | Up | 0.118 | 6.31 | 7.00 | $4.85 \times 10^{-11}$ | $5.66 \times 10^{-9}$ |
| *DENND4C* | Up | 0.070 | 6.45 | 7.00 | $4.86 \times 10^{-11}$ | $5.66 \times 10^{-9}$ |
| *EEF2K* | Up | 0.091 | 4.83 | 6.99 | $5.09 \times 10^{-11}$ | $5.88 \times 10^{-9}$ |
| *PRDM5* | Up | 0.101 | 4.52 | 6.99 | $5.13 \times 10^{-11}$ | $5.90 \times 10^{-9}$ |
| *MNS1* | Up | 0.091 | 4.37 | 6.99 | $5.18 \times 10^{-11}$ | $5.94 \times 10^{-9}$ |
| *SMAD5* | Up | 0.076 | 4.83 | 6.98 | $5.34 \times 10^{-11}$ | $6.09 \times 10^{-9}$ |
| *TTC39C* | Up | 0.089 | 7.12 | 6.97 | $5.76 \times 10^{-11}$ | $6.47 \times 10^{-9}$ |
| ENSG00000261829 | Up | 0.207 | 1.39 | 6.97 | $5.93 \times 10^{-11}$ | $6.64 \times 10^{-9}$ |
| *SCX* | Up | 0.109 | 2.13 | 6.95 | $6.48 \times 10^{-11}$ | $7.19 \times 10^{-9}$ |
| *PXN* | Up | 0.111 | 2.28 | 6.91 | $7.98 \times 10^{-11}$ | $8.80 \times 10^{-9}$ |
| *PDK4-AS1* | Up | 0.165 | 2.17 | 6.90 | $8.34 \times 10^{-11}$ | $9.12 \times 10^{-9}$ |
| ENSG00000260296 | Up | 0.168 | 0.84 | 6.90 | $8.58 \times 10^{-11}$ | $9.34 \times 10^{-9}$ |
| *FAM161A* | Up | 0.093 | 4.13 | 6.89 | $9.00 \times 10^{-11}$ | $9.73 \times 10^{-9}$ |
| *ITGA2B* | Up | 0.105 | 2.69 | 6.89 | $9.10 \times 10^{-11}$ | $9.79 \times 10^{-9}$ |
| *RSAD2* | Up | 0.093 | 3.71 | 6.89 | $9.19 \times 10^{-11}$ | $9.85 \times 10^{-9}$ |
| *UNG* | Up | 0.082 | 2.67 | 6.89 | $9.22 \times 10^{-11}$ | $9.85 \times 10^{-9}$ |
| *TFAP4* | Up | 0.084 | 2.54 | 6.88 | $9.68 \times 10^{-11}$ | $1.02 \times 10^{-8}$ |
| *BTN3A1* | Up | 0.190 | 0.58 | 6.88 | $9.71 \times 10^{-11}$ | $1.02 \times 10^{-8}$ |
| *ZBTB16* | Up | 0.107 | 8.21 | 6.88 | $9.74 \times 10^{-11}$ | $1.02 \times 10^{-8}$ |
| *LRRC17* | Up | 0.080 | 3.07 | 6.87 | $9.92 \times 10^{-11}$ | $1.04 \times 10^{-8}$ |
| *KCNC4* | Up | 0.081 | 4.32 | 6.87 | $1.01 \times 10^{-10}$ | $1.05 \times 10^{-8}$ |
| *MAMDC2* | Up | 0.122 | 5.39 | 6.87 | $1.01 \times 10^{-10}$ | $1.05 \times 10^{-8}$ |
| *KIF13B* | Up | 0.116 | 5.21 | 6.86 | $1.07 \times 10^{-10}$ | $1.10 \times 10^{-8}$ |
| *IL15* | Up | 0.229 | 1.26 | 6.85 | $1.15 \times 10^{-10}$ | $1.18 \times 10^{-8}$ |
| *IMPA1* | Up | 0.099 | 4.46 | 6.84 | $1.22 \times 10^{-10}$ | $1.24 \times 10^{-8}$ |
| *ROPN1L* | Up | 0.103 | 2.28 | 6.82 | $1.33 \times 10^{-10}$ | $1.34 \times 10^{-8}$ |
| *LPL* | Up | 0.188 | 2.15 | 6.81 | $1.38 \times 10^{-10}$ | $1.38 \times 10^{-8}$ |
| *HBP1* | Up | 0.069 | 5.31 | 6.80 | $1.47 \times 10^{-10}$ | $1.46 \times 10^{-8}$ |
| *RESF1* | Up | 0.078 | 5.10 | 6.79 | $1.54 \times 10^{-10}$ | $1.52 \times 10^{-8}$ |
| *FBXO40* | Up | 0.142 | 2.46 | 6.79 | $1.54 \times 10^{-10}$ | $1.52 \times 10^{-8}$ |
| *CA3* | Up | 0.145 | 4.36 | 6.79 | $1.56 \times 10^{-10}$ | $1.53 \times 10^{-8}$ |
| *HABP2* | Up | 0.180 | 1.28 | 6.78 | $1.66 \times 10^{-10}$ | $1.63 \times 10^{-8}$ |
| *WASF2* | Up | 0.073 | 4.73 | 6.77 | $1.75 \times 10^{-10}$ | $1.70 \times 10^{-8}$ |
| *SLC17A5* | Up | 0.068 | 5.60 | 6.76 | $1.84 \times 10^{-10}$ | $1.78 \times 10^{-8}$ |
| *PSMC2* | Up | 0.050 | 4.87 | 6.76 | $1.87 \times 10^{-10}$ | $1.81 \times 10^{-8}$ |
| *KIF7* | Up | 0.105 | 2.25 | 6.76 | $1.89 \times 10^{-10}$ | $1.82 \times 10^{-8}$ |
| *TGFA* | Up | 0.133 | 1.15 | 6.75 | $1.96 \times 10^{-10}$ | $1.88 \times 10^{-8}$ |
| *SLC18B1* | Up | 0.094 | 4.70 | 6.74 | $2.02 \times 10^{-10}$ | $1.93 \times 10^{-8}$ |
| *NAALAD2* | Up | 0.083 | 6.08 | 6.74 | $2.10 \times 10^{-10}$ | $1.99 \times 10^{-8}$ |
| *EPHX2* | Up | 0.120 | 1.65 | 6.74 | $2.11 \times 10^{-10}$ | $1.99 \times 10^{-8}$ |
| *LINC02359* | Up | 0.188 | 2.63 | 6.73 | $2.21 \times 10^{-10}$ | $2.08 \times 10^{-8}$ |
| *GINM1* | Up | 0.067 | 4.71 | 6.72 | $2.27 \times 10^{-10}$ | $2.13 \times 10^{-8}$ |

| Gene | Direction with age | log2 fold change per decade | Average log2 CPM | Moderated t | P value | FDR |
|---|---|---|---|---|---|---|
| *TMEM237* | Up | 0.078 | 3.89 | 6.72 | $2.32 \times 10^{-10}$ | $2.16 \times 10^{-8}$ |
| *UBXN2A* | Up | 0.071 | 5.01 | 6.72 | $2.33 \times 10^{-10}$ | $2.17 \times 10^{-8}$ |
| *TOR1AIP1* | Up | 0.061 | 4.10 | 6.71 | $2.45 \times 10^{-10}$ | $2.25 \times 10^{-8}$ |
| *COTL1* | Up | 0.098 | 2.54 | 6.70 | $2.57 \times 10^{-10}$ | $2.34 \times 10^{-8}$ |
| *SEC14L6* | Up | 0.151 | 0.52 | 6.68 | $2.83 \times 10^{-10}$ | $2.55 \times 10^{-8}$ |
| *EPCIP-AS1* | Up | 0.115 | 3.63 | 6.68 | $2.94 \times 10^{-10}$ | $2.64 \times 10^{-8}$ |
| *NHS* | Up | 0.104 | 5.25 | 6.67 | $3.01 \times 10^{-10}$ | $2.69 \times 10^{-8}$ |
| *CCDC92* | Up | 0.088 | 6.86 | 6.67 | $3.07 \times 10^{-10}$ | $2.73 \times 10^{-8}$ |
| *DAPK1* | Up | 0.098 | 7.70 | 6.66 | $3.23 \times 10^{-10}$ | $2.87 \times 10^{-8}$ |
| *ACACB* | Up | 0.100 | 2.74 | 6.65 | $3.31 \times 10^{-10}$ | $2.93 \times 10^{-8}$ |
| ENSG00000258465 | Up | 0.117 | 3.11 | 6.65 | $3.35 \times 10^{-10}$ | $2.96 \times 10^{-8}$ |
| *MTHFD2L* | Up | 0.120 | 5.66 | 6.64 | $3.55 \times 10^{-10}$ | $3.10 \times 10^{-8}$ |
| *TLE2* | Up | 0.103 | 3.20 | 6.63 | $3.77 \times 10^{-10}$ | $3.27 \times 10^{-8}$ |
| *PAN2* | Up | 0.067 | 3.61 | 6.63 | $3.82 \times 10^{-10}$ | $3.31 \times 10^{-8}$ |
| *ALDH6A1* | Up | 0.072 | 3.06 | 6.63 | $3.88 \times 10^{-10}$ | $3.34 \times 10^{-8}$ |
| *LMAN1* | Up | 0.082 | 4.31 | 6.62 | $4.01 \times 10^{-10}$ | $3.43 \times 10^{-8}$ |
| *NCAPG2* | Up | 0.076 | 3.86 | 6.62 | $4.04 \times 10^{-10}$ | $3.44 \times 10^{-8}$ |
| *NNT-AS1* | Up | 0.105 | 4.55 | 6.61 | $4.15 \times 10^{-10}$ | $3.52 \times 10^{-8}$ |
| *KBTBD12* | Up | 0.093 | 5.43 | 6.61 | $4.23 \times 10^{-10}$ | $3.58 \times 10^{-8}$ |
| *OSCAR* | Up | 0.178 | 1.50 | 6.61 | $4.33 \times 10^{-10}$ | $3.65 \times 10^{-8}$ |
| *MIOS* | Up | 0.070 | 4.97 | 6.61 | $4.33 \times 10^{-10}$ | $3.65 \times 10^{-8}$ |
| ENSG00000225498 | Up | 0.125 | 2.50 | 6.60 | $4.38 \times 10^{-10}$ | $3.67 \times 10^{-8}$ |
| *RPA1* | Up | 0.075 | 4.41 | 6.60 | $4.42 \times 10^{-10}$ | $3.69 \times 10^{-8}$ |
| *GRK6* | Up | 0.078 | 3.26 | 6.60 | $4.54 \times 10^{-10}$ | $3.77 \times 10^{-8}$ |
| *GPR55* | Up | 0.223 | 1.36 | 6.60 | $4.56 \times 10^{-10}$ | $3.78 \times 10^{-8}$ |
| ENSG00000261543 | Up | 0.256 | 2.24 | 6.59 | $4.61 \times 10^{-10}$ | $3.81 \times 10^{-8}$ |
| *RBBP9* | Up | 0.088 | 1.96 | 6.59 | $4.78 \times 10^{-10}$ | $3.93 \times 10^{-8}$ |
| *NACC2* | Up | 0.108 | 1.38 | 6.58 | $4.88 \times 10^{-10}$ | $3.99 \times 10^{-8}$ |
| *RET* | Up | 0.145 | 1.40 | 6.58 | $4.90 \times 10^{-10}$ | $3.99 \times 10^{-8}$ |
| *ANKRD13A* | Up | 0.088 | 4.33 | 6.58 | $5.00 \times 10^{-10}$ | $4.05 \times 10^{-8}$ |
| *DUBR* | Up | 0.105 | 6.73 | 6.58 | $5.04 \times 10^{-10}$ | $4.07 \times 10^{-8}$ |
| *MSRB3* | Up | 0.101 | 5.67 | 6.57 | $5.16 \times 10^{-10}$ | $4.14 \times 10^{-8}$ |
| *TNS3* | Up | 0.138 | 2.39 | 6.57 | $5.23 \times 10^{-10}$ | $4.18 \times 10^{-8}$ |
| *MAP1LC3B2* | Up | 0.135 | 2.25 | 6.57 | $5.31 \times 10^{-10}$ | $4.20 \times 10^{-8}$ |
| *TMEM145* | Up | 0.073 | 4.00 | 6.56 | $5.42 \times 10^{-10}$ | $4.27 \times 10^{-8}$ |
| *PAK2* | Up | 0.062 | 5.52 | 6.56 | $5.42 \times 10^{-10}$ | $4.27 \times 10^{-8}$ |
| ENSG00000236829 | Up | 0.091 | 1.77 | 6.56 | $5.43 \times 10^{-10}$ | $4.27 \times 10^{-8}$ |
| *PHKA2* | Up | 0.080 | 4.42 | 6.56 | $5.45 \times 10^{-10}$ | $4.27 \times 10^{-8}$ |
| *ZKSCAN8* | Up | 0.086 | 4.13 | 6.55 | $5.69 \times 10^{-10}$ | $4.45 \times 10^{-8}$ |
| *VASH1-DT* | Up | 0.099 | 3.01 | 6.51 | $7.40 \times 10^{-10}$ | $5.65 \times 10^{-8}$ |
| ENSG00000253108 | Up | 0.180 | 0.07 | 6.50 | $7.50 \times 10^{-10}$ | $5.72 \times 10^{-8}$ |
| ENSG00000228877 | Up | 0.186 | 0.46 | 6.50 | $7.57 \times 10^{-10}$ | $5.75 \times 10^{-8}$ |
| *SPARCL1* | Up | 0.116 | 8.07 | 6.50 | $7.58 \times 10^{-10}$ | $5.75 \times 10^{-8}$ |
| *SGPP2* | Up | 0.072 | 5.66 | 6.50 | $7.68 \times 10^{-10}$ | $5.79 \times 10^{-8}$ |
| *BAIAP3* | Up | 0.108 | 4.74 | 6.50 | $7.72 \times 10^{-10}$ | $5.80 \times 10^{-8}$ |
| *NIPSNAP3B* | Up | 0.090 | 3.61 | 6.50 | $7.74 \times 10^{-10}$ | $5.81 \times 10^{-8}$ |
| *LINC02033* | Up | 0.106 | 0.23 | 6.49 | $8.03 \times 10^{-10}$ | $6.01 \times 10^{-8}$ |
| *LINC02054* | Up | 0.118 | 2.75 | 6.48 | $8.72 \times 10^{-10}$ | $6.45 \times 10^{-8}$ |
| ENSG00000280107 | Up | 0.191 | 0.46 | 6.48 | $8.73 \times 10^{-10}$ | $6.45 \times 10^{-8}$ |
| *RBL1* | Up | 0.062 | 4.30 | 6.47 | $8.77 \times 10^{-10}$ | $6.45 \times 10^{-8}$ |
| *IL12A-AS1* | Up | 0.143 | 5.30 | 6.47 | $9.00 \times 10^{-10}$ | $6.58 \times 10^{-8}$ |
| *RNF216* | Up | 0.068 | 6.41 | 6.46 | $9.41 \times 10^{-10}$ | $6.86 \times 10^{-8}$ |
| *ATAD2* | Up | 0.082 | 5.36 | 6.45 | $9.89 \times 10^{-10}$ | $7.17 \times 10^{-8}$ |
| *KIF27* | Up | 0.055 | 4.85 | 6.44 | $1.04 \times 10^{-9}$ | $7.48 \times 10^{-8}$ |
| ENSG00000205794 | Up | 0.148 | 0.29 | 6.42 | $1.16 \times 10^{-9}$ | $8.30 \times 10^{-8}$ |
| *MBP* | Up | 0.091 | 6.33 | 6.42 | $1.17 \times 10^{-9}$ | $8.33 \times 10^{-8}$ |
| *LURAP1L* | Up | 0.093 | 3.75 | 6.41 | $1.24 \times 10^{-9}$ | $8.77 \times 10^{-8}$ |

| Gene | Direction with age | log2 fold change per decade | Average log2 CPM | Moderated t | P value | FDR |
|---|---|---|---|---|---|---|
| *RUNX1-AS1* | Up | 0.181 | 1.42 | 6.41 | $1.27 \times 10^{-9}$ | $9.00 \times 10^{-8}$ |
| *TTC28* | Up | 0.074 | 7.86 | 6.39 | $1.35 \times 10^{-9}$ | $9.50 \times 10^{-8}$ |
| ENSG00000260664 | Up | 0.167 | 5.09 | 6.39 | $1.37 \times 10^{-9}$ | $9.59 \times 10^{-8}$ |
| *SMCR8* | Up | 0.093 | 2.78 | 6.39 | $1.39 \times 10^{-9}$ | $9.73 \times 10^{-8}$ |
| *DPY19L4* | Up | 0.060 | 4.20 | 6.38 | $1.47 \times 10^{-9}$ | $1.02 \times 10^{-7}$ |
| *VASH2* | Up | 0.142 | 2.27 | 6.37 | $1.52 \times 10^{-9}$ | $1.05 \times 10^{-7}$ |
| *AMPH* | Up | 0.084 | 8.22 | 6.37 | $1.54 \times 10^{-9}$ | $1.06 \times 10^{-7}$ |
| *KDM6B* | Up | 0.091 | 3.10 | 6.37 | $1.56 \times 10^{-9}$ | $1.07 \times 10^{-7}$ |
| *CNTN2* | Up | 0.081 | 4.18 | 6.36 | $1.58 \times 10^{-9}$ | $1.08 \times 10^{-7}$ |
| ENSG00000225032 | Up | 0.109 | 0.69 | 6.36 | $1.61 \times 10^{-9}$ | $1.09 \times 10^{-7}$ |
| *ACP6* | Up | 0.099 | 3.59 | 6.36 | $1.61 \times 10^{-9}$ | $1.10 \times 10^{-7}$ |
| *CERKL* | Up | 0.168 | 3.13 | 6.36 | $1.63 \times 10^{-9}$ | $1.10 \times 10^{-7}$ |
| *FAM178B* | Up | 0.110 | 3.60 | 6.35 | $1.68 \times 10^{-9}$ | $1.13 \times 10^{-7}$ |
| *HMGN5* | Up | 0.117 | 3.91 | 6.34 | $1.80 \times 10^{-9}$ | $1.21 \times 10^{-7}$ |
| *TSPYL2* | Up | 0.076 | 6.15 | 6.34 | $1.82 \times 10^{-9}$ | $1.22 \times 10^{-7}$ |
| ENSG00000260418 | Up | 0.124 | 2.32 | 6.34 | $1.82 \times 10^{-9}$ | $1.22 \times 10^{-7}$ |
| ENSG00000287458 | Up | 0.301 | –2.62 | 6.33 | $1.90 \times 10^{-9}$ | $1.26 \times 10^{-7}$ |
| ENSG00000286891 | Up | 0.153 | 3.15 | 6.33 | $1.91 \times 10^{-9}$ | $1.26 \times 10^{-7}$ |
| *PLEKHF1* | Up | 0.186 | –1.60 | 6.33 | $1.91 \times 10^{-9}$ | $1.26 \times 10^{-7}$ |
| ENSG00000273118 | Up | 0.177 | 4.65 | 6.32 | $2.01 \times 10^{-9}$ | $1.33 \times 10^{-7}$ |
| *SH3D19* | Up | 0.067 | 6.63 | 6.32 | $2.04 \times 10^{-9}$ | $1.34 \times 10^{-7}$ |
| *APOL2* | Up | 0.115 | 4.19 | 6.31 | $2.11 \times 10^{-9}$ | $1.38 \times 10^{-7}$ |
| *GSTM4* | Up | 0.098 | 1.97 | 6.30 | $2.23 \times 10^{-9}$ | $1.45 \times 10^{-7}$ |
| *RTKN2* | Up | 0.101 | 4.40 | 6.30 | $2.26 \times 10^{-9}$ | $1.46 \times 10^{-7}$ |
| *FRYL* | Up | 0.077 | 7.08 | 6.30 | $2.27 \times 10^{-9}$ | $1.46 \times 10^{-7}$ |
| *AK3* | Up | 0.100 | 4.45 | 6.30 | $2.27 \times 10^{-9}$ | $1.46 \times 10^{-7}$ |
| *ANKRD35* | Up | 0.083 | 2.59 | 6.30 | $2.27 \times 10^{-9}$ | $1.46 \times 10^{-7}$ |
| *TRMT11* | Up | 0.073 | 8.09 | 6.29 | $2.39 \times 10^{-9}$ | $1.53 \times 10^{-7}$ |
| *CLU* | Up | 0.138 | 6.45 | 6.29 | $2.40 \times 10^{-9}$ | $1.53 \times 10^{-7}$ |
| *EPN3* | Up | 0.186 | –0.92 | 6.28 | $2.41 \times 10^{-9}$ | $1.53 \times 10^{-7}$ |
| *NREP* | Down | –0.212 | 6.83 | –13.43 | $6.12 \times 10^{-29}$ | $5.80 \times 10^{-25}$ |
| *DMBT1L1* | Down | –0.440 | 1.21 | –12.58 | $1.99 \times 10^{-26}$ | $1.41 \times 10^{-22}$ |
| *MKRN3* | Down | –0.286 | 1.95 | –12.50 | $3.21 \times 10^{-26}$ | $1.83 \times 10^{-22}$ |
| *ADAMTS9* | Down | –0.350 | 2.73 | –12.33 | $1.06 \times 10^{-25}$ | $3.77 \times 10^{-22}$ |
| ENSG00000237250 | Down | –0.394 | 1.91 | –11.68 | $8.24 \times 10^{-24}$ | $2.60 \times 10^{-20}$ |
| *UACA* | Down | –0.232 | 1.52 | –11.01 | $7.22 \times 10^{-22}$ | $1.61 \times 10^{-18}$ |
| *C6orf141* | Down | –0.521 | 1.07 | –11.01 | $7.38 \times 10^{-22}$ | $1.61 \times 10^{-18}$ |
| *OTOF* | Down | –0.271 | 1.46 | –10.82 | $2.54 \times 10^{-21}$ | $4.81 \times 10^{-18}$ |
| *WNT6* | Down | –0.330 | –0.39 | –10.60 | $1.08 \times 10^{-20}$ | $1.91 \times 10^{-17}$ |
| *ARHGAP10* | Down | –0.239 | 6.06 | –10.58 | $1.24 \times 10^{-20}$ | $2.07 \times 10^{-17}$ |
| *ELFN1* | Down | –0.269 | 0.39 | –10.30 | $7.43 \times 10^{-20}$ | $1.10 \times 10^{-16}$ |
| *STUM* | Down | –0.229 | 5.24 | –10.30 | $7.54 \times 10^{-20}$ | $1.10 \times 10^{-16}$ |
| *HIP1* | Down | –0.210 | 4.44 | –10.30 | $7.72 \times 10^{-20}$ | $1.10 \times 10^{-16}$ |
| *NREP-AS1* | Down | –0.467 | 2.39 | –10.15 | $2.01 \times 10^{-19}$ | $2.59 \times 10^{-16}$ |
| *ADAMTS9-AS1* | Down | –0.413 | 1.25 | –10.13 | $2.35 \times 10^{-19}$ | $2.78 \times 10^{-16}$ |
| *ALPL* | Down | –0.304 | 1.44 | –10.01 | $5.11 \times 10^{-19}$ | $5.58 \times 10^{-16}$ |
| *SOX4* | Down | –0.261 | 2.16 | –9.96 | $7.05 \times 10^{-19}$ | $7.42 \times 10^{-16}$ |
| ENSG00000271860 | Down | –0.202 | 7.30 | –9.89 | $1.10 \times 10^{-18}$ | $1.08 \times 10^{-15}$ |
| *RASGRP1* | Down | –0.133 | 6.06 | –9.75 | $2.68 \times 10^{-18}$ | $2.54 \times 10^{-15}$ |
| *EXPH5* | Down | –0.248 | 3.93 | –9.57 | $8.82 \times 10^{-18}$ | $7.84 \times 10^{-15}$ |
| *LINC01509* | Down | –0.286 | –0.03 | –9.47 | $1.60 \times 10^{-17}$ | $1.38 \times 10^{-14}$ |
| ENSG00000287251 | Down | –0.387 | –1.25 | –9.22 | $7.86 \times 10^{-17}$ | $5.73 \times 10^{-14}$ |
| *P2RY6* | Down | –0.333 | 1.13 | –9.20 | $8.80 \times 10^{-17}$ | $6.25 \times 10^{-14}$ |
| *STARD4-AS1* | Down | –0.244 | 6.31 | –9.14 | $1.36 \times 10^{-16}$ | $9.27 \times 10^{-14}$ |
| *LINC02197* | Down | –0.250 | 3.73 | –9.14 | $1.37 \times 10^{-16}$ | $9.27 \times 10^{-14}$ |
| *COL24A1* | Down | –0.221 | 6.72 | –9.10 | $1.69 \times 10^{-16}$ | $1.12 \times 10^{-13}$ |
| *LINC01052* | Down | –0.443 | –1.89 | –9.07 | $2.06 \times 10^{-16}$ | $1.27 \times 10^{-13}$ |

| Gene | Direction with age | log2 fold change per decade | Average log2 CPM | Moderated t | P value | FDR |
|---|---|---|---|---|---|---|
| ENSG00000287373 | Down | –0.350 | 4.02 | –9.06 | $2.15 \times 10^{-16}$ | $1.30 \times 10^{-13}$ |
| *HCRTR1* | Down | –0.334 | –0.03 | –9.04 | $2.50 \times 10^{-16}$ | $1.48 \times 10^{-13}$ |
| *MYO16* | Down | –0.179 | 5.09 | –8.99 | $3.33 \times 10^{-16}$ | $1.86 \times 10^{-13}$ |
| ENSG00000259072 | Down | –0.269 | 3.82 | –8.97 | $3.94 \times 10^{-16}$ | $2.14 \times 10^{-13}$ |
| *DSG3* | Down | –0.381 | –0.15 | –8.94 | $4.55 \times 10^{-16}$ | $2.39 \times 10^{-13}$ |
| *ZNF420* | Down | –0.074 | 5.72 | –8.93 | $4.99 \times 10^{-16}$ | $2.55 \times 10^{-13}$ |
| *LINC02717* | Down | –0.419 | 1.15 | –8.93 | $5.02 \times 10^{-16}$ | $2.55 \times 10^{-13}$ |
| *CACNA1G* | Down | –0.137 | 4.54 | –8.64 | $2.93 \times 10^{-15}$ | $1.30 \times 10^{-12}$ |
| *STARD4* | Down | –0.142 | 4.76 | –8.56 | $4.89 \times 10^{-15}$ | $2.11 \times 10^{-12}$ |
| *LUZP2* | Down | –0.202 | 6.49 | –8.52 | $6.37 \times 10^{-15}$ | $2.62 \times 10^{-12}$ |
| *LINC02725* | Down | –0.216 | 1.15 | –8.48 | $7.81 \times 10^{-15}$ | $3.13 \times 10^{-12}$ |
| *LINC02133* | Down | –0.191 | 2.24 | –8.46 | $9.04 \times 10^{-15}$ | $3.57 \times 10^{-12}$ |
| *CACNA1H* | Down | –0.244 | 1.32 | –8.44 | $1.04 \times 10^{-14}$ | $4.00 \times 10^{-12}$ |
| *CFAP20DC* | Down | –0.187 | 4.97 | –8.41 | $1.21 \times 10^{-14}$ | $4.54 \times 10^{-12}$ |
| *MPL* | Down | –0.315 | 0.74 | –8.38 | $1.51 \times 10^{-14}$ | $5.52 \times 10^{-12}$ |
| ENSG00000226699 | Down | –0.265 | –0.48 | –8.35 | $1.79 \times 10^{-14}$ | $6.31 \times 10^{-12}$ |
| *ST8SIA2* | Down | –0.411 | 0.18 | –8.35 | $1.80 \times 10^{-14}$ | $6.31 \times 10^{-12}$ |
| ENSG00000258696 | Down | –0.233 | 0.22 | –8.29 | $2.60 \times 10^{-14}$ | $8.49 \times 10^{-12}$ |
| *HDC* | Down | –0.273 | –0.96 | –8.23 | $3.70 \times 10^{-14}$ | $1.19 \times 10^{-11}$ |
| ENSG00000258107 | Down | –0.219 | 3.32 | –8.22 | $3.96 \times 10^{-14}$ | $1.25 \times 10^{-11}$ |
| *C12orf42* | Down | –0.187 | 3.82 | –8.21 | $4.06 \times 10^{-14}$ | $1.27 \times 10^{-11}$ |
| *OR9Q1* | Down | –0.172 | 2.06 | –8.17 | $5.38 \times 10^{-14}$ | $1.61 \times 10^{-11}$ |
| *RAMP3* | Down | –0.191 | 1.89 | –8.17 | $5.39 \times 10^{-14}$ | $1.61 \times 10^{-11}$ |
| *CAMK1* | Down | –0.101 | 4.88 | –8.13 | $6.71 \times 10^{-14}$ | $1.97 \times 10^{-11}$ |
| ENSG00000277332 | Down | –0.342 | 0.15 | –8.06 | $1.05 \times 10^{-13}$ | $3.05 \times 10^{-11}$ |
| *CFAP20DC-AS1* | Down | –0.148 | 5.47 | –8.01 | $1.38 \times 10^{-13}$ | $3.97 \times 10^{-11}$ |
| *APLF* | Down | –0.133 | 4.45 | –8.00 | $1.47 \times 10^{-13}$ | $4.15 \times 10^{-11}$ |
| *TMC1* | Down | –0.151 | 1.94 | –8.00 | $1.48 \times 10^{-13}$ | $4.15 \times 10^{-11}$ |
| *FOXO6* | Down | –0.202 | 0.51 | –7.96 | $1.84 \times 10^{-13}$ | $4.88 \times 10^{-11}$ |
| *KIF21B* | Down | –0.136 | 4.76 | –7.95 | $2.02 \times 10^{-13}$ | $5.26 \times 10^{-11}$ |
| *SLC1A6* | Down | –0.115 | 2.95 | –7.94 | $2.08 \times 10^{-13}$ | $5.37 \times 10^{-11}$ |
| *HES4* | Down | –0.183 | 3.43 | –7.94 | $2.14 \times 10^{-13}$ | $5.43 \times 10^{-11}$ |
| *ADRA1D* | Down | –0.246 | –0.07 | –7.93 | $2.20 \times 10^{-13}$ | $5.52 \times 10^{-11}$ |
| *NETO2* | Down | –0.112 | 6.40 | –7.89 | $2.85 \times 10^{-13}$ | $6.93 \times 10^{-11}$ |
| *LINC02237* | Down | –0.301 | 2.23 | –7.88 | $3.01 \times 10^{-13}$ | $7.19 \times 10^{-11}$ |
| *INF2* | Down | –0.130 | 4.56 | –7.83 | $4.08 \times 10^{-13}$ | $9.28 \times 10^{-11}$ |
| *IQANK1* | Down | –0.134 | 3.40 | –7.80 | $4.88 \times 10^{-13}$ | $1.09 \times 10^{-10}$ |
| *RERGL* | Down | –0.216 | 2.45 | –7.79 | $5.19 \times 10^{-13}$ | $1.15 \times 10^{-10}$ |
| *LINC01965* | Down | –0.185 | 3.56 | –7.76 | $6.16 \times 10^{-13}$ | $1.35 \times 10^{-10}$ |
| *GPR26* | Down | –0.172 | 3.73 | –7.75 | $6.43 \times 10^{-13}$ | $1.39 \times 10^{-10}$ |
| *ABCC3* | Down | –0.143 | 2.33 | –7.74 | $6.95 \times 10^{-13}$ | $1.48 \times 10^{-10}$ |
| *TIE1* | Down | –0.283 | 0.26 | –7.72 | $7.64 \times 10^{-13}$ | $1.61 \times 10^{-10}$ |
| *JCAD* | Down | –0.142 | 5.49 | –7.71 | $8.16 \times 10^{-13}$ | $1.70 \times 10^{-10}$ |
| ENSG00000286222 | Down | –0.316 | –0.83 | –7.69 | $9.43 \times 10^{-13}$ | $1.96 \times 10^{-10}$ |
| ENSG00000286321 | Down | –0.342 | 0.73 | –7.68 | $9.79 \times 10^{-13}$ | $2.02 \times 10^{-10}$ |
| ENSG00000259048 | Down | –0.174 | 5.20 | –7.67 | $1.07 \times 10^{-12}$ | $2.19 \times 10^{-10}$ |
| *ADAMTS15* | Down | –0.213 | –0.97 | –7.63 | $1.29 \times 10^{-12}$ | $2.54 \times 10^{-10}$ |
| *DOK6* | Down | –0.139 | 8.19 | –7.63 | $1.34 \times 10^{-12}$ | $2.59 \times 10^{-10}$ |
| *DLGAP1-AS4* | Down | –0.090 | 7.24 | –7.62 | $1.44 \times 10^{-12}$ | $2.76 \times 10^{-10}$ |
| *TTC9B* | Down | –0.162 | 4.45 | –7.61 | $1.51 \times 10^{-12}$ | $2.89 \times 10^{-10}$ |
| *MEIS3* | Down | –0.117 | 5.59 | –7.57 | $1.83 \times 10^{-12}$ | $3.40 \times 10^{-10}$ |
| *FREM3* | Down | –0.187 | 4.71 | –7.54 | $2.22 \times 10^{-12}$ | $4.05 \times 10^{-10}$ |
| *LIMS2* | Down | –0.194 | 0.84 | –7.51 | $2.70 \times 10^{-12}$ | $4.82 \times 10^{-10}$ |
| *LINC02572* | Down | –0.148 | 1.44 | –7.49 | $2.91 \times 10^{-12}$ | $5.14 \times 10^{-10}$ |
| *PRKG2* | Down | –0.153 | 4.91 | –7.49 | $2.97 \times 10^{-12}$ | $5.21 \times 10^{-10}$ |
| ENSG00000286738 | Down | –0.165 | 3.14 | –7.44 | $3.93 \times 10^{-12}$ | $6.80 \times 10^{-10}$ |
| *STAC* | Down | –0.199 | 0.41 | –7.44 | $3.95 \times 10^{-12}$ | $6.80 \times 10^{-10}$ |

| Gene | Direction with age | log2 fold change per decade | Average log2 CPM | Moderated t | P value | FDR |
|---|---|---|---|---|---|---|
| ENSG00000286003 | Down | –0.218 | 2.87 | –7.41 | $4.67 \times 10^{-12}$ | $7.95 \times 10^{-10}$ |
| ENSG00000234464 | Down | –0.227 | 3.16 | –7.38 | $5.81 \times 10^{-12}$ | $9.72 \times 10^{-10}$ |
| *PNPLA3* | Down | –0.203 | 1.44 | –7.36 | $6.17 \times 10^{-12}$ | $1.02 \times 10^{-9}$ |
| *LINC01205* | Down | –0.181 | 2.10 | –7.36 | $6.28 \times 10^{-12}$ | $1.03 \times 10^{-9}$ |
| *HES1* | Down | –0.227 | 0.59 | –7.36 | $6.38 \times 10^{-12}$ | $1.04 \times 10^{-9}$ |
| ENSG00000243276 | Down | –0.140 | 3.56 | –7.34 | $7.09 \times 10^{-12}$ | $1.14 \times 10^{-9}$ |
| ENSG00000242029 | Down | –0.186 | 2.38 | –7.34 | $7.21 \times 10^{-12}$ | $1.15 \times 10^{-9}$ |
| *ITGB5* | Down | –0.138 | 1.30 | –7.32 | $7.83 \times 10^{-12}$ | $1.23 \times 10^{-9}$ |
| *TTC7A* | Down | –0.076 | 5.15 | –7.29 | $9.35 \times 10^{-12}$ | $1.43 \times 10^{-9}$ |
| *LINC02873* | Down | –0.304 | –2.33 | –7.29 | $9.55 \times 10^{-12}$ | $1.44 \times 10^{-9}$ |
| *RYR3* | Down | –0.170 | 7.16 | –7.29 | $9.75 \times 10^{-12}$ | $1.47 \times 10^{-9}$ |
| *PLCH2* | Down | –0.146 | 3.75 | –7.27 | $1.04 \times 10^{-11}$ | $1.55 \times 10^{-9}$ |
| ENSG00000285634 | Down | –0.198 | 4.89 | –7.27 | $1.05 \times 10^{-11}$ | $1.56 \times 10^{-9}$ |
| *CACNA1S* | Down | –0.168 | 0.32 | –7.26 | $1.10 \times 10^{-11}$ | $1.61 \times 10^{-9}$ |
| *ARHGAP20* | Down | –0.083 | 6.71 | –7.25 | $1.17 \times 10^{-11}$ | $1.69 \times 10^{-9}$ |
| *NEURL1B* | Down | –0.131 | 3.33 | –7.25 | $1.18 \times 10^{-11}$ | $1.69 \times 10^{-9}$ |
| *KCNMB2* | Down | –0.354 | 3.92 | –7.25 | $1.22 \times 10^{-11}$ | $1.74 \times 10^{-9}$ |
| *LINC02280* | Down | –0.248 | –0.64 | –7.23 | $1.32 \times 10^{-11}$ | $1.84 \times 10^{-9}$ |
| *TWIST2* | Down | –0.221 | 2.95 | –7.23 | $1.34 \times 10^{-11}$ | $1.85 \times 10^{-9}$ |
| *ZNF354C* | Down | –0.104 | 3.29 | –7.21 | $1.53 \times 10^{-11}$ | $2.09 \times 10^{-9}$ |
| *TRAF5* | Down | –0.138 | 3.84 | –7.18 | $1.74 \times 10^{-11}$ | $2.35 \times 10^{-9}$ |
| *RASL10A* | Down | –0.156 | 4.07 | –7.18 | $1.80 \times 10^{-11}$ | $2.40 \times 10^{-9}$ |
| ENSG00000287567 | Down | –0.254 | 0.74 | –7.17 | $1.83 \times 10^{-11}$ | $2.43 \times 10^{-9}$ |
| *LINC01208* | Down | –0.135 | 3.79 | –7.17 | $1.84 \times 10^{-11}$ | $2.43 \times 10^{-9}$ |
| *COL16A1* | Down | –0.094 | 3.91 | –7.17 | $1.88 \times 10^{-11}$ | $2.47 \times 10^{-9}$ |
| ENSG00000286141 | Down | –0.298 | 0.50 | –7.14 | $2.18 \times 10^{-11}$ | $2.84 \times 10^{-9}$ |
| *OTUB2* | Down | –0.138 | 1.48 | –7.14 | $2.19 \times 10^{-11}$ | $2.85 \times 10^{-9}$ |
| ENSG00000287038 | Down | –0.201 | 1.48 | –7.13 | $2.33 \times 10^{-11}$ | $2.99 \times 10^{-9}$ |
| *AGBL1-AS1* | Down | –0.362 | –1.37 | –7.13 | $2.42 \times 10^{-11}$ | $3.09 \times 10^{-9}$ |
| *COL22A1* | Down | –0.264 | 2.50 | –7.11 | $2.66 \times 10^{-11}$ | $3.33 \times 10^{-9}$ |
| ENSG00000262267 | Down | –0.188 | 2.86 | –7.08 | $3.17 \times 10^{-11}$ | $3.87 \times 10^{-9}$ |
| *FGF1* | Down | –0.285 | 4.85 | –7.06 | $3.59 \times 10^{-11}$ | $4.36 \times 10^{-9}$ |
| *NTM-AS1* | Down | –0.215 | 2.25 | –7.05 | $3.80 \times 10^{-11}$ | $4.60 \times 10^{-9}$ |
| *ZFP37* | Down | –0.110 | 3.73 | –7.02 | $4.27 \times 10^{-11}$ | $5.09 \times 10^{-9}$ |
| ENSG00000285980 | Down | –0.242 | 3.08 | –7.02 | $4.35 \times 10^{-11}$ | $5.17 \times 10^{-9}$ |
| *PLCZ1* | Down | –0.311 | –0.65 | –7.02 | $4.36 \times 10^{-11}$ | $5.17 \times 10^{-9}$ |
| *CD226* | Down | –0.133 | 3.70 | –7.02 | $4.40 \times 10^{-11}$ | $5.19 \times 10^{-9}$ |
| ENSG00000236920 | Down | –0.230 | –0.51 | –7.00 | $4.99 \times 10^{-11}$ | $5.79 \times 10^{-9}$ |
| *LINC01333* | Down | –0.286 | 1.18 | –6.99 | $5.36 \times 10^{-11}$ | $6.09 \times 10^{-9}$ |
| *B3GNT2* | Down | –0.085 | 3.77 | –6.98 | $5.61 \times 10^{-11}$ | $6.35 \times 10^{-9}$ |
| *PAFAH1B2P2* | Down | –0.125 | 2.59 | –6.97 | $5.76 \times 10^{-11}$ | $6.47 \times 10^{-9}$ |
| *SSBP4* | Down | –0.080 | 5.29 | –6.96 | $6.23 \times 10^{-11}$ | $6.95 \times 10^{-9}$ |
| ENSG00000230730 | Down | –0.163 | 0.25 | –6.95 | $6.56 \times 10^{-11}$ | $7.26 \times 10^{-9}$ |
| *ZNF350* | Down | –0.114 | 2.93 | –6.91 | $8.26 \times 10^{-11}$ | $9.07 \times 10^{-9}$ |
| *SYT17* | Down | –0.134 | 5.00 | –6.89 | $8.92 \times 10^{-11}$ | $9.68 \times 10^{-9}$ |
| *LINC02543* | Down | –0.331 | –1.59 | –6.88 | $9.32 \times 10^{-11}$ | $9.93 \times 10^{-9}$ |
| *MFSD10* | Down | –0.101 | 3.76 | –6.88 | $9.55 \times 10^{-11}$ | $1.01 \times 10^{-8}$ |
| *LINC01141* | Down | –0.138 | 2.69 | –6.87 | $1.01 \times 10^{-10}$ | $1.05 \times 10^{-8}$ |
| *DNAJB5* | Down | –0.204 | 1.91 | –6.85 | $1.14 \times 10^{-10}$ | $1.17 \times 10^{-8}$ |
| ENSG00000287072 | Down | –0.321 | 1.13 | –6.85 | $1.15 \times 10^{-10}$ | $1.17 \times 10^{-8}$ |
| *TRPC4* | Down | –0.116 | 4.85 | –6.83 | $1.23 \times 10^{-10}$ | $1.25 \times 10^{-8}$ |
| *PMCHL2* | Down | –0.349 | –1.43 | –6.82 | $1.33 \times 10^{-10}$ | $1.34 \times 10^{-8}$ |
| ENSG00000238117 | Down | –0.185 | 1.43 | –6.81 | $1.45 \times 10^{-10}$ | $1.44 \times 10^{-8}$ |
| *HMGCS1* | Down | –0.115 | 5.27 | –6.80 | $1.52 \times 10^{-10}$ | $1.50 \times 10^{-8}$ |
| ENSG00000279286 | Down | –0.193 | –0.08 | –6.78 | $1.69 \times 10^{-10}$ | $1.64 \times 10^{-8}$ |
| *AGBL1* | Down | –0.180 | 4.92 | –6.74 | $2.06 \times 10^{-10}$ | $1.96 \times 10^{-8}$ |
| *HAS2-AS1* | Down | –0.174 | 2.39 | –6.72 | $2.30 \times 10^{-10}$ | $2.15 \times 10^{-8}$ |

| Gene | Direction with age | log2 fold change per decade | Average log2 CPM | Moderated t | P value | FDR |
|---|---|---|---|---|---|---|
| *ZNF350-AS1* | Down | –0.135 | 3.63 | –6.71 | $2.40 \times 10^{-10}$ | $2.22 \times 10^{-8}$ |
| *PLCD3* | Down | –0.116 | 0.96 | –6.71 | $2.40 \times 10^{-10}$ | $2.22 \times 10^{-8}$ |
| ENSG00000237654 | Down | –0.211 | 3.23 | –6.71 | $2.41 \times 10^{-10}$ | $2.22 \times 10^{-8}$ |
| ENSG00000288036 | Down | –0.176 | 5.66 | –6.71 | $2.50 \times 10^{-10}$ | $2.28 \times 10^{-8}$ |
| *TPRG1* | Down | –0.173 | 3.18 | –6.69 | $2.74 \times 10^{-10}$ | $2.49 \times 10^{-8}$ |
| *LINC02462* | Down | –0.239 | 0.13 | –6.69 | $2.76 \times 10^{-10}$ | $2.49 \times 10^{-8}$ |
| *GRM2* | Down | –0.113 | 3.38 | –6.68 | $2.83 \times 10^{-10}$ | $2.55 \times 10^{-8}$ |
| ENSG00000227958 | Down | –0.264 | 1.66 | –6.65 | $3.36 \times 10^{-10}$ | $2.96 \times 10^{-8}$ |
| ENSG00000231698 | Down | –0.188 | –0.38 | –6.64 | $3.54 \times 10^{-10}$ | $3.10 \times 10^{-8}$ |
| *RN7SL814P* | Down | –0.297 | –0.41 | –6.64 | $3.54 \times 10^{-10}$ | $3.10 \times 10^{-8}$ |
| *TECTA* | Down | –0.100 | 3.30 | –6.63 | $3.88 \times 10^{-10}$ | $3.34 \times 10^{-8}$ |
| ENSG00000227407 | Down | –0.142 | 2.24 | –6.62 | $3.92 \times 10^{-10}$ | $3.36 \times 10^{-8}$ |
| *MAN1A2P1* | Down | –0.209 | –0.63 | –6.62 | $3.92 \times 10^{-10}$ | $3.36 \times 10^{-8}$ |
| *ZFP69* | Down | –0.129 | 1.76 | –6.60 | $4.42 \times 10^{-10}$ | $3.69 \times 10^{-8}$ |
| ENSG00000259035 | Down | –0.264 | 2.72 | –6.59 | $4.66 \times 10^{-10}$ | $3.84 \times 10^{-8}$ |
| *LINC02144* | Down | –0.138 | 1.70 | –6.59 | $4.79 \times 10^{-10}$ | $3.93 \times 10^{-8}$ |
| *TCF7L1* | Down | –0.126 | 2.07 | –6.58 | $4.96 \times 10^{-10}$ | $4.03 \times 10^{-8}$ |
| ENSG00000256196 | Down | –0.127 | 1.46 | –6.57 | $5.11 \times 10^{-10}$ | $4.12 \times 10^{-8}$ |
| *NKD1* | Down | –0.171 | 0.42 | –6.57 | $5.19 \times 10^{-10}$ | $4.16 \times 10^{-8}$ |
| *JAM2* | Down | –0.117 | 4.81 | –6.57 | $5.27 \times 10^{-10}$ | $4.19 \times 10^{-8}$ |
| ENSG00000251377 | Down | –0.172 | 0.94 | –6.57 | $5.28 \times 10^{-10}$ | $4.19 \times 10^{-8}$ |
| *FBLN2* | Down | –0.194 | 0.91 | –6.55 | $5.86 \times 10^{-10}$ | $4.56 \times 10^{-8}$ |
| ENSG00000226207 | Down | –0.311 | –1.29 | –6.52 | $6.78 \times 10^{-10}$ | $5.27 \times 10^{-8}$ |
| ENSG00000248431 | Down | –0.230 | 4.38 | –6.52 | $6.83 \times 10^{-10}$ | $5.28 \times 10^{-8}$ |
| *MGC32805* | Down | –0.244 | 1.14 | –6.52 | $6.84 \times 10^{-10}$ | $5.28 \times 10^{-8}$ |
| *TAMALIN* | Down | –0.106 | 4.16 | –6.52 | $6.91 \times 10^{-10}$ | $5.32 \times 10^{-8}$ |
| *PIK3C2G* | Down | –0.170 | 3.54 | –6.52 | $6.99 \times 10^{-10}$ | $5.37 \times 10^{-8}$ |
| ENSG00000287844 | Down | –0.160 | 3.10 | –6.51 | $7.27 \times 10^{-10}$ | $5.57 \times 10^{-8}$ |
| *TTYH1* | Down | –0.081 | 4.93 | –6.50 | $7.63 \times 10^{-10}$ | $5.77 \times 10^{-8}$ |
| *MYLIP* | Down | –0.105 | 3.62 | –6.49 | $8.05 \times 10^{-10}$ | $6.01 \times 10^{-8}$ |
| *PRKG2-AS1* | Down | –0.255 | 1.77 | –6.49 | $8.12 \times 10^{-10}$ | $6.04 \times 10^{-8}$ |
| ENSG00000258649 | Down | –0.227 | –0.24 | –6.48 | $8.46 \times 10^{-10}$ | $6.28 \times 10^{-8}$ |
| *ZNF90* | Down | –0.118 | 1.27 | –6.47 | $8.78 \times 10^{-10}$ | $6.45 \times 10^{-8}$ |
| ENSG00000250706 | Down | –0.146 | 2.31 | –6.47 | $8.95 \times 10^{-10}$ | $6.55 \times 10^{-8}$ |
| *AGRN* | Down | –0.139 | 2.05 | –6.45 | $9.84 \times 10^{-10}$ | $7.15 \times 10^{-8}$ |
| *CYP26B1* | Down | –0.152 | 2.25 | –6.44 | $1.04 \times 10^{-9}$ | $7.48 \times 10^{-8}$ |
| *FAUP4* | Down | –0.122 | 1.98 | –6.44 | $1.04 \times 10^{-9}$ | $7.48 \times 10^{-8}$ |
| *PTPRB* | Down | –0.081 | 6.36 | –6.44 | $1.07 \times 10^{-9}$ | $7.70 \times 10^{-8}$ |
| *DOCK7-DT* | Down | –0.121 | 1.88 | –6.42 | $1.17 \times 10^{-9}$ | $8.33 \times 10^{-8}$ |
| *OGG1* | Down | –0.093 | 4.52 | –6.42 | $1.18 \times 10^{-9}$ | $8.38 \times 10^{-8}$ |
| ENSG00000265702 | Down | –0.139 | 1.28 | –6.40 | $1.33 \times 10^{-9}$ | $9.40 \times 10^{-8}$ |
| *CALML3-AS1* | Down | –0.154 | 0.00 | –6.39 | $1.40 \times 10^{-9}$ | $9.78 \times 10^{-8}$ |
| ENSG00000225649 | Down | –0.183 | 4.33 | –6.39 | $1.42 \times 10^{-9}$ | $9.87 \times 10^{-8}$ |
| ENSG00000288102 | Down | –0.294 | 5.63 | –6.37 | $1.54 \times 10^{-9}$ | $1.06 \times 10^{-7}$ |
| *SCN3B* | Down | –0.067 | 6.48 | –6.36 | $1.58 \times 10^{-9}$ | $1.08 \times 10^{-7}$ |
| *GRIN3A* | Down | –0.152 | 4.32 | –6.36 | $1.59 \times 10^{-9}$ | $1.08 \times 10^{-7}$ |
| *HOMER1* | Down | –0.126 | 8.06 | –6.36 | $1.63 \times 10^{-9}$ | $1.10 \times 10^{-7}$ |
| *XKR4* | Down | –0.088 | 8.79 | –6.36 | $1.65 \times 10^{-9}$ | $1.11 \times 10^{-7}$ |
| *PLPPR1* | Down | –0.120 | 4.66 | –6.33 | $1.88 \times 10^{-9}$ | $1.25 \times 10^{-7}$ |
| *FGD3* | Down | –0.097 | 0.48 | –6.33 | $1.89 \times 10^{-9}$ | $1.26 \times 10^{-7}$ |
| ENSG00000223930 | Down | –0.241 | 5.54 | –6.31 | $2.07 \times 10^{-9}$ | $1.36 \times 10^{-7}$ |
| *KLF2-DT* | Down | –0.122 | 3.07 | –6.31 | $2.08 \times 10^{-9}$ | $1.36 \times 10^{-7}$ |
| *KREMEN2* | Down | –0.207 | 0.26 | –6.31 | $2.11 \times 10^{-9}$ | $1.38 \times 10^{-7}$ |
| ENSG00000263823 | Down | –0.174 | 0.47 | –6.30 | $2.19 \times 10^{-9}$ | $1.43 \times 10^{-7}$ |
| *GYPE* | Down | –0.192 | 3.58 | –6.30 | $2.29 \times 10^{-9}$ | $1.47 \times 10^{-7}$ |
| ENSG00000261810 | Down | –0.125 | 3.02 | –6.28 | $2.50 \times 10^{-9}$ | $1.59 \times 10^{-7}$ |
| ENSG00000287887 | Down | –0.159 | –0.24 | –6.27 | $2.55 \times 10^{-9}$ | $1.61 \times 10^{-7}$ |

| Gene | Direction with age | log2 fold change per decade | Average log2 CPM | Moderated t | P value | FDR |
|---|---|---|---|---|---|---|
| *USP12* | Down | –0.063 | 5.59 | –6.27 | $2.61 \times 10^{-9}$ | $1.65 \times 10^{-7}$ |
| *TTC6* | Down | –0.180 | 5.79 | –6.27 | $2.65 \times 10^{-9}$ | $1.66 \times 10^{-7}$ |
| *LINGO3* | Down | –0.178 | –1.20 | –6.26 | $2.68 \times 10^{-9}$ | $1.68 \times 10^{-7}$ |
| *MIR3667HG* | Down | –0.151 | 1.73 | –6.26 | $2.80 \times 10^{-9}$ | $1.73 \times 10^{-7}$ |
| *LINC02008* | Down | –0.183 | 3.46 | –6.25 | $2.93 \times 10^{-9}$ | $1.80 \times 10^{-7}$ |
| ENSG00000250493 | Down | –0.213 | 2.25 | –6.25 | $2.97 \times 10^{-9}$ | $1.82 \times 10^{-7}$ |
| *NFRKB* | Down | –0.059 | 3.80 | –6.24 | $2.97 \times 10^{-9}$ | $1.82 \times 10^{-7}$ |
| *LINC00378* | Down | –0.167 | 3.82 | –6.23 | $3.28 \times 10^{-9}$ | $1.97 \times 10^{-7}$ |
| *NPHS1* | Down | –0.125 | 1.92 | –6.22 | $3.39 \times 10^{-9}$ | $2.03 \times 10^{-7}$ |
| *NFIB-AS1* | Down | –0.143 | 1.63 | –6.22 | $3.39 \times 10^{-9}$ | $2.03 \times 10^{-7}$ |
| *TMEM215* | Down | –0.290 | –0.74 | –6.21 | $3.51 \times 10^{-9}$ | $2.09 \times 10^{-7}$ |
| ENSG00000227863 | Down | –0.117 | 3.22 | –6.21 | $3.62 \times 10^{-9}$ | $2.16 \times 10^{-7}$ |
| *MAPK15* | Down | –0.229 | –0.73 | –6.20 | $3.74 \times 10^{-9}$ | $2.21 \times 10^{-7}$ |
| *GPR160* | Down | –0.141 | 1.94 | –6.19 | $3.99 \times 10^{-9}$ | $2.34 \times 10^{-7}$ |
| *ENPP7P15* | Down | –0.128 | 3.24 | –6.18 | $4.13 \times 10^{-9}$ | $2.41 \times 10^{-7}$ |
| ENSG00000287203 | Down | –0.323 | –2.54 | –6.17 | $4.30 \times 10^{-9}$ | $2.50 \times 10^{-7}$ |
| *MYLK* | Down | –0.101 | 5.09 | –6.17 | $4.48 \times 10^{-9}$ | $2.59 \times 10^{-7}$ |
| ENSG00000269296 | Down | –0.280 | –1.36 | –6.17 | $4.51 \times 10^{-9}$ | $2.61 \times 10^{-7}$ |
| *TP53INP2* | Down | –0.081 | 1.75 | –6.16 | $4.62 \times 10^{-9}$ | $2.66 \times 10^{-7}$ |
| *PTGER3* | Down | –0.292 | 2.48 | –6.16 | $4.65 \times 10^{-9}$ | $2.67 \times 10^{-7}$ |
| *LECT2* | Down | –0.346 | –1.04 | –6.15 | $5.01 \times 10^{-9}$ | $2.87 \times 10^{-7}$ |
| *GPC6-AS1* | Down | –0.264 | 1.84 | –6.15 | $5.03 \times 10^{-9}$ | $2.87 \times 10^{-7}$ |
| ENSG00000226939 | Down | –0.183 | 1.15 | –6.14 | $5.16 \times 10^{-9}$ | $2.93 \times 10^{-7}$ |
| *SPEG* | Down | –0.075 | 4.31 | –6.11 | $6.01 \times 10^{-9}$ | $3.36 \times 10^{-7}$ |
| *BDNF* | Down | –0.142 | 4.34 | –6.11 | $6.10 \times 10^{-9}$ | $3.39 \times 10^{-7}$ |
| *PAPSS2* | Down | –0.128 | 4.78 | –6.11 | $6.16 \times 10^{-9}$ | $3.42 \times 10^{-7}$ |
| *MAPT-AS1* | Down | –0.068 | 4.30 | –6.10 | $6.18 \times 10^{-9}$ | $3.43 \times 10^{-7}$ |
| *ASB18* | Down | –0.205 | 1.60 | –6.10 | $6.25 \times 10^{-9}$ | $3.45 \times 10^{-7}$ |
| ENSG00000255367 | Down | –0.130 | 3.63 | –6.10 | $6.25 \times 10^{-9}$ | $3.45 \times 10^{-7}$ |
| *DSG2* | Down | –0.165 | 0.59 | –6.08 | $7.02 \times 10^{-9}$ | $3.82 \times 10^{-7}$ |
| ENSG00000265643 | Down | –0.269 | 5.01 | –6.08 | $7.05 \times 10^{-9}$ | $3.82 \times 10^{-7}$ |
| *SFTA1P* | Down | –0.242 | 3.86 | –6.08 | $7.08 \times 10^{-9}$ | $3.82 \times 10^{-7}$ |
| *SORCS2* | Down | –0.114 | 6.40 | –6.07 | $7.39 \times 10^{-9}$ | $3.96 \times 10^{-7}$ |
| *WSB1* | Down | –0.081 | 7.36 | –6.07 | $7.53 \times 10^{-9}$ | $4.02 \times 10^{-7}$ |
| *NOPCHAP1* | Down | –0.070 | 5.02 | –6.07 | $7.55 \times 10^{-9}$ | $4.03 \times 10^{-7}$ |
| *PROM1* | Down | –0.125 | 0.39 | –6.07 | $7.57 \times 10^{-9}$ | $4.03 \times 10^{-7}$ |
| *TRIM46* | Down | –0.067 | 3.99 | –6.06 | $7.87 \times 10^{-9}$ | $4.18 \times 10^{-7}$ |
| *ZNF271P* | Down | –0.063 | 4.69 | –6.04 | $8.47 \times 10^{-9}$ | $4.46 \times 10^{-7}$ |
| *KIF2B* | Down | –0.303 | –1.22 | –6.04 | $8.75 \times 10^{-9}$ | $4.59 \times 10^{-7}$ |
| *ZNF571* | Down | –0.059 | 4.07 | –6.02 | $9.54 \times 10^{-9}$ | $4.95 \times 10^{-7}$ |
| *NEK2* | Down | –0.205 | –0.73 | –6.02 | $9.59 \times 10^{-9}$ | $4.95 \times 10^{-7}$ |
| *DLK2* | Down | –0.098 | 2.21 | –6.02 | $9.59 \times 10^{-9}$ | $4.95 \times 10^{-7}$ |
| *CELSR1* | Down | –0.117 | 1.74 | –6.02 | $9.67 \times 10^{-9}$ | $4.98 \times 10^{-7}$ |
| *ZNF883* | Down | –0.105 | 5.37 | –6.01 | $9.95 \times 10^{-9}$ | $5.11 \times 10^{-7}$ |
| ENSG00000266840 | Down | –0.243 | 1.42 | –6.01 | $9.97 \times 10^{-9}$ | $5.11 \times 10^{-7}$ |
| *KIF26A* | Down | –0.147 | –0.41 | –6.01 | $9.98 \times 10^{-9}$ | $5.11 \times 10^{-7}$ |
| *NSMCE1-DT* | Down | –0.098 | 2.46 | –6.01 | $1.01 \times 10^{-8}$ | $5.14 \times 10^{-7}$ |
| *BDNF-AS* | Down | –0.095 | 4.92 | –6.01 | $1.03 \times 10^{-8}$ | $5.22 \times 10^{-7}$ |
| ENSG00000284703 | Down | –0.183 | 0.61 | –6.00 | $1.05 \times 10^{-8}$ | $5.33 \times 10^{-7}$ |
| *RMEL3* | Down | –0.124 | 1.58 | –6.00 | $1.05 \times 10^{-8}$ | $5.33 \times 10^{-7}$ |
| *KIRREL2* | Down | –0.177 | –0.97 | –6.00 | $1.07 \times 10^{-8}$ | $5.37 \times 10^{-7}$ |
| *LINC01501* | Down | –0.129 | 1.22 | –5.99 | $1.14 \times 10^{-8}$ | $5.64 \times 10^{-7}$ |
